# Virtual Environments for Research into Social Evolution (VERSE): A novel experimental environment for the study of human social learning

**DOI:** 10.1101/2022.08.24.505099

**Authors:** C. Easter

## Abstract

Social learning (learning from others) can be a cost-effective way of gaining information compared to asocial (independent) learning. However, learning from others indiscriminately can lead to the acquisition of maladaptive behaviours or outdated information. Evolutionary theory therefore predicts that individuals will use social information adaptively through the use of ‘social learning strategies’. Restrictive laboratory conditions, however, make studying human learning strategies problematic. Abstract tasks, unrealistic sources of social information and methodologies that do not take into account the influence of physical location over large spaces make it difficult to ascertain if previous findings are representative of the way we would use social information in reality. Here I describe a novel platform for studying human social behaviour within immersive virtual environments: “Virtual Environments for Research into Social Evolution” (VERSE). Through the use of gaming technology, VERSE allows researchers to build realistic, three-dimensional, open world environments where participants can complete ecologically relevant tasks while actively observing computer-controlled artificial intelligence agents (AIs) that act as realistic yet controllable sources of social information. This methodological article begins by exploring what social learning strategies are and the problems with studying social learning behaviour in humans (compared to animal populations, for example). I then discuss how gaming technology can be used in behavioural research and follow on with a detailed account of the specific functionalities available in VERSE. I conclude with a worked example of how VERSE can be used to construct a novel behavioural experiment. Altogether, VERSE has great potential to give us insight into how human individuals learn within novel environments in a way that has never before been possible.

## 2. Background

Social learning (learning from others) occurs in many species and can be a highly cost-effective way of gaining information (Hoppitt and Laland, 2013), making it adaptive in many biological contexts (Galef and Laland, 2005) over independent, or ‘asocial’, learning. There are many ways in which individuals can gain information from one another – from direct observation of an individual to interactions with the products of its behaviour – and many specific mechanisms of social learning – from ‘local enhancement’, where a demonstrator’s behaviour attracts an observer’s attention to a particular stimulus, to direct imitation of a specific behaviour or set of behaviours (Heyes, 1994). Research suggests that both animals and humans are also selective in the way they use social information, deploying ‘social learning strategies’ that dictate when, what and from whom they learn socially (Laland, 2004; Rendell *et al*., 2011). The use of such selective social information use is expected to be adaptive compared to copying individuals indiscriminately (Boyd and Richerson, 1988; Schlag, 1998; Rendell *et al*., 2010), as a constant lack of direct sampling from the environment can lead to the spread of maladaptive or outdated behaviours through the population (Laland and Williams, 1998; Rogers, 1988). Evidence suggests that both humans and non-human animals are capable of using social information adaptively and that there are similarities in the use of social learning strategies between different taxa.

However, making direct comparisons between the way humans and animals use social information is difficult, not least because of the hugely different methodologies used to study humans versus non-human animals. Animal experiments generally involve individuals learning ecologically relevant survival skills (e.g. foraging, predator avoidance) through the observations of informed demonstrators during either traditional ‘demonstrator-observer’ experiments or larger scale, more naturalistic ‘open diffusion’ experiments. Take, for example, Allen and colleagues (2013) study on the spread of the ‘lobtail feeding’ behaviour through a population of humpback whales (*Megaptera novaeangliae*) in the Gulf of Maine, which used specialised, network-based statistical analyses to investigate natural patterns of information transmission through freely interacting social groups. In contrast, most human experiments lack realism for three major reasons.

Firstly, they are often limited by highly abstract tasks with little ecological relevance that offer little in the way of behavioural flexibility, e.g. choosing between coloured options (Efferson *et al.,* 2008; Toelch *et al.,* 2010), deciding which of two lines is the longest (Morgan *et al.,* 2012), or building towers using modelling clay and spaghetti (Caldwell and Eve, 2014). Such tasks often have a limited number of arbitrary solutions and thus offer little in the way of behavioural flexibility or scope for participants to exploit prior knowledge or skills – which I argue could inflate participants’ reliance on social information simply to avoid guesswork. Secondly, social information is provided to participants in unrealistic ways, e.g. as flashing tiles (Morgan *et al.,* 2012) or numerical values showing the frequency or success of behaviours being used in the population (Toelch *et al*., 2010; Mesoudi and O’Brien, 2008). This arguably oversimplifies the social learning process – participants are effectively being told what other individuals are doing and how useful their behaviours are, while in reality, we would need to actively observe others, process this information before deciding which elements (if any) to copy. Thirdly, while animal experiments have the potential to span huge home ranges, human experiments are almost always restricted to the confines of the lab, or some other relatively small indoor space (e.g. Reader *et al*., 2008; Whiten *et al.,* 2016). In other words, human experiments are extremely spatially limited, with tasks taking place in very localised areas. In reality, many tasks associated with human survival (e.g. hunter-gatherer activities) take place over kilometre scales (Hamilton *et al*., 2007; Whallon, 2006). Such spatial scales are simply not possible using traditional lab-based methods, but are likely to have a substantial influence over who is able to learn from whom at a particular time (particularly if demonstrated behaviours occur asynchronously over time and space) (Coussi-Korbel and Fragaszy, 1995), as well as the social learning mechanisms that are adopted (e.g. spatial scale may have a more pronounced effect on imitation, which requires close observation, than local enhancement, which requires only an attraction to a particular location due to the presence of conspecifics) (Heyes, 1994). Overall, this means that current methodologies struggle to consider the ecological and evolutionary foundations of human social behaviour.

One largely unexploited methodology that may allow humans to be studied in more ‘natural’, spatially realistic conditions, and in a framework more comparable to field experiments on animals, is virtual reality (VR). Computer-based experiments, in general, offer a greater possibility of extending their reach to the general public, and hence a more diverse pool of participants, than lab-based experiments (Vicens *et al*., 2018). Relatively simple computer-based tasks have already proved useful in the field for the creation of novel tasks (e.g. Mesoudi 2008; Mesoudi and O’Brien, 2008; Morgan *et al*., 2012). Mesoudi and colleagues’ research in particular demonstrates how virtual tasks can be created that would represent real-life challenges in traditional human communities – in this case, designing virtual arrowheads to use in a virtual hunting ground (Mesoudi 2008; Mesoudi and O’Brien, 2008). Such computer-based methodologies have even extended to the study of human cumulative culture – as in Miu and colleagues’ (2018) study of collaborative computer programming. Perhaps most importantly for the study of human behavioural ecology and social evolution, VR also gives us a unique opportunity to study human behaviour in survival situations that cannot be replicated under experimental conditions, e.g. the outbreak of a fire (Arias *et al.,* 2018) or evacuation scenarios (Moussaïd *et al.,* 2016).

Large, open world, three-dimensional environments that participants can navigate freely have the potential to allow realistic social interactions between networks of individuals across realistic spatial scales, without the need to leave the lab. Massively multiplayer online role-playing games (MMORPGs) such as *World of Warcraft* offer a particularly exciting opportunity to study real world social dynamics within VR. This potential was highlighted by a virtual ‘disease outbreak’ that plagued *World of Warcraft* in 2005 (Lofgren and Fefferman, 2007; Balicer, 2007). Due to the realistic movement patterns and social interactions of players, the disease was able to spread through the virtual population in a way analogous to real world disease dynamics. The virtual outbreak also gave insights into human behavioural responses to unexpected events – something that is notoriously difficult to model due to the complexity and unpredictability of human nature. MMORPGs could also act as experimental environments for studying the development of cultural norms across different ‘societies’ (Strimling and Frey, 2020). However, despite the immense potential of commercial multiplayer games for the study of human social evolution, these games are not purpose built for behavioural experiments. Ideally, social learning researchers could benefit from a VR platform built specifically to study human social behaviour within realistic, ecologically relevant environments. Developing complex, realistic virtual worlds for behavioural research is not a simple task and requires game coding expertise not readily available to most researchers – hence the full potential of VR for studying human social learning within realistic three-dimensional spaces has not been fully exploited.

Here, I describe a novel tool specifically developed for the purpose of studying human social learning using virtual reality: “Virtual Environments for Research into Social Evolution” (VERSE). Developed using Unity3D game development technology, VERSE gives researchers the ability to create complex, immersive 3D environments containing ecologically relevant tasks and challenges, without the need for game coding knowledge. Within VERSE environments, participants take control of a virtual player to explore and learn within realistic 3D spaces. Computer-controlled artificial intelligence agents (AIs), programmed by the researcher to behave in a certain way, offer optional sources of social information. VERSE is designed with the limitations of laboratory-based human social learning experiments in mind and offers a novel way for human behaviour to be studied in ‘wild’ environments. In particular, VERSE is designed to give participants the freedom to navigate 3D environments; allow tasks to span large spatial scales; provide more realistic sources of social information, in the sense that participants must actively observe an individual’s behaviour and decide how to use this information; allow humans to be studied in naturalistic environments and survival scenarios; expose participants to realistic, ecologically-relevant tasks that can optionally require cumulative behaviours to complete; and provide researchers with a flexible toolkit that can be used to create potentially infinite new environments, tasks and types of social information due to its modular design. VERSE, therefore, has great potential for human social learning research, allowing humans to be studied in a comparable framework to animal research, allowing the replication of animal studies using human subjects and giving us a glimpse into how our social behaviour may have aided us in our evolutionary past.

What follows is a detailed account of VERSE, including the features already available in Unity3D that make it a suitable program for this type of research and features created specifically for VERSE to provide researchers with the ability to generate their own complex social learning environments.

## 3. Unity3D in Behavioural Research

Unity3D^1^ is one of a series of virtual reality packages that have been used to create realistic environments for use as educational tools (e.g. Houghton *et al*., 2015) and systems for studying human behaviour (e.g. Arias *et al.,* 2018). Unity3D boasts a range of features that make it suitable for use within research projects on social evolution. Here, I outline some of these key features. Table 4.1 describes some of the main Unity3D terminology intended to aid the reading of this document.

**Table 0.1.**
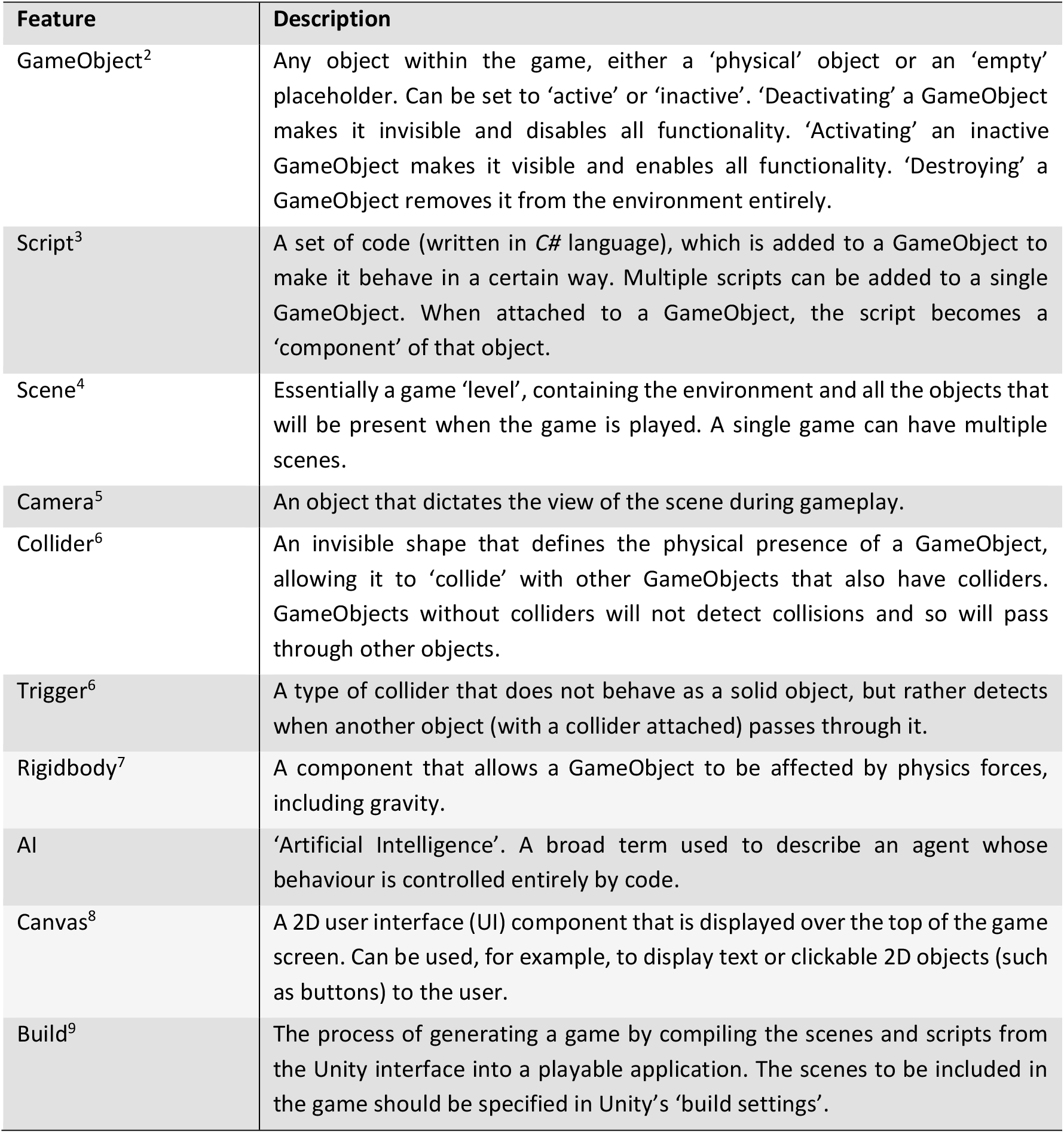
Descriptions of some of the Unity terminology referred to in this document.

| Feature | Description |
| --- | --- |
| GameObject<sup>2</sup> | Any object within the game, either a 'physical' object or an 'empty' placeholder. Can be set to 'active' or 'inactive'. 'Deactivating' a GameObject makes it invisible and disables all functionality. 'Activating' an inactive GameObject makes it visible and enables all functionality. 'Destroying' a GameObject removes it from the environment entirely. |
| Script<sup>3</sup> | A set of code (written in C# language), which is added to a GameObject to make it behave in a certain way. Multiple scripts can be added to a single GameObject. When attached to a GameObject, the script becomes a 'component' of that object. |
| Scene<sup>4</sup> | Essentially a game 'level', containing the environment and all the objects that will be present when the game is played. A single game can have multiple scenes. |
| Camera<sup>5</sup> | An object that dictates the view of the scene during gameplay. |
| Collider<sup>6</sup> | An invisible shape that defines the physical presence of a GameObject, allowing it to 'collide' with other GameObjects that also have colliders. GameObjects without colliders will not detect collisions and so will pass through other objects. |
| Trigger<sup>6</sup> | A type of collider that does not behave as a solid object, but rather detects when another object (with a collider attached) passes through it. |
| Rigidbody<sup>7</sup> | A component that allows a GameObject to be affected by physics forces, including gravity. |
| AI | 'Artificial Intelligence'. A broad term used to describe an agent whose behaviour is controlled entirely by code. |
| Canvas<sup>8</sup> | A 2D user interface (UI) component that is displayed over the top of the game screen. Can be used, for example, to display text or clickable 2D objects (such as buttons) to the user. |
| Build<sup>9</sup> | The process of generating a game by compiling the scenes and scripts from the Unity interface into a playable application. The scenes to be included in the game should be specified in Unity's 'build settings'. |

### 3.1. Flexible coding, ScriptableObjects and inheritance

Unity3D uses *C#* game coding which offers a high level of flexibility for the creation of tasks and character behaviours. Each section of code is contained within a script and so can be attached to different GameObjects in different combinations, thus allowing scripts to act as ‘building blocks’ for more complex character behaviour and task functionalities. Data containers called ‘ScriptableObjects’ can also be used to store particular properties within the Unity interface, allowing them to be shared between different scripts or objects during gameplay^10^. ScriptableObjects are notably used within VERSE for applying modifications to ‘interactable objects’ (section 4.4.3) and creating ‘weather conditions’ within the environment (section 4.4.7.1). In addition, Unity’s ‘inheritance’ system^11^ is particularly useful for creating ‘parent’ scripts which contain a number of functions and properties that are shared with (or ‘inherited’ by) any number of ‘child’ scripts. Thus, one can easily create multiple scripts of a particular type with shared functionality without having to repeat blocks of code. For example, inheritance is used within VERSE to create ‘interactable objects’ of various different types (section 4.4.3).

### 3.2. The Asset Store

Unity3D’s Asset Store^12^ allows game developers to obtain additional features for their games, such as 3D models or specialised scripts. VERSE uses one feature from the Asset Store, a third-person character controller (see below), that was modified for purpose and is included within the VERSE system. VERSE is an inclusive package, containing all the features and coding required for researchers to generate their own social learning environments, and so the Asset Store is not essential for its use. However, researchers can benefit from the Asset Store, for example, if they wish to obtain 3D models to use in tasks or as other features within the environment.

### 3.3. Environment creation

Creating VERSE in Unity means that researchers can take advantage of Unity3D’s built-in terrain tool^13^, which is particularly helpful if a researcher is wishing to conduct (or replicate) a study within a naturalistic environment. The terrain tool allows the easy creation of realistic environments with varied topology and allows the rapid addition of trees, vegetation and surface textures (Figure 4.1). This tool can also be used to convert real Ordnance Survey 2D map data to 3D virtual terrains (Robinson *et al.,* 2015). Terrains can potentially be any size – the creation of large terrains being of particular use if a researcher wishes to allow their participants to have free movement over large scales. Hunter-gatherer processes, for example, occur over kilometre scales (90km^2^ to 730,000km^2^ according to Hamilton *et al*., 2007). Such scales are not possible in conventional lab experiments, but become relatively easy to achieve, subject to computational demands, in virtual reality. Terrains can also be generated that vary in their complexity, allowing hypotheses concerning the effect of environmental complexity on social information use to be investigated. Such hypotheses have been tested in animals – for example, Webster *et al*. (2013) demonstrated that threespine sticklebacks (*Gasterosteus aculeatus*) were more likely to use social information in structured environments – but current methodologies make comparative studies on humans difficult.

**Figure 0.1.**
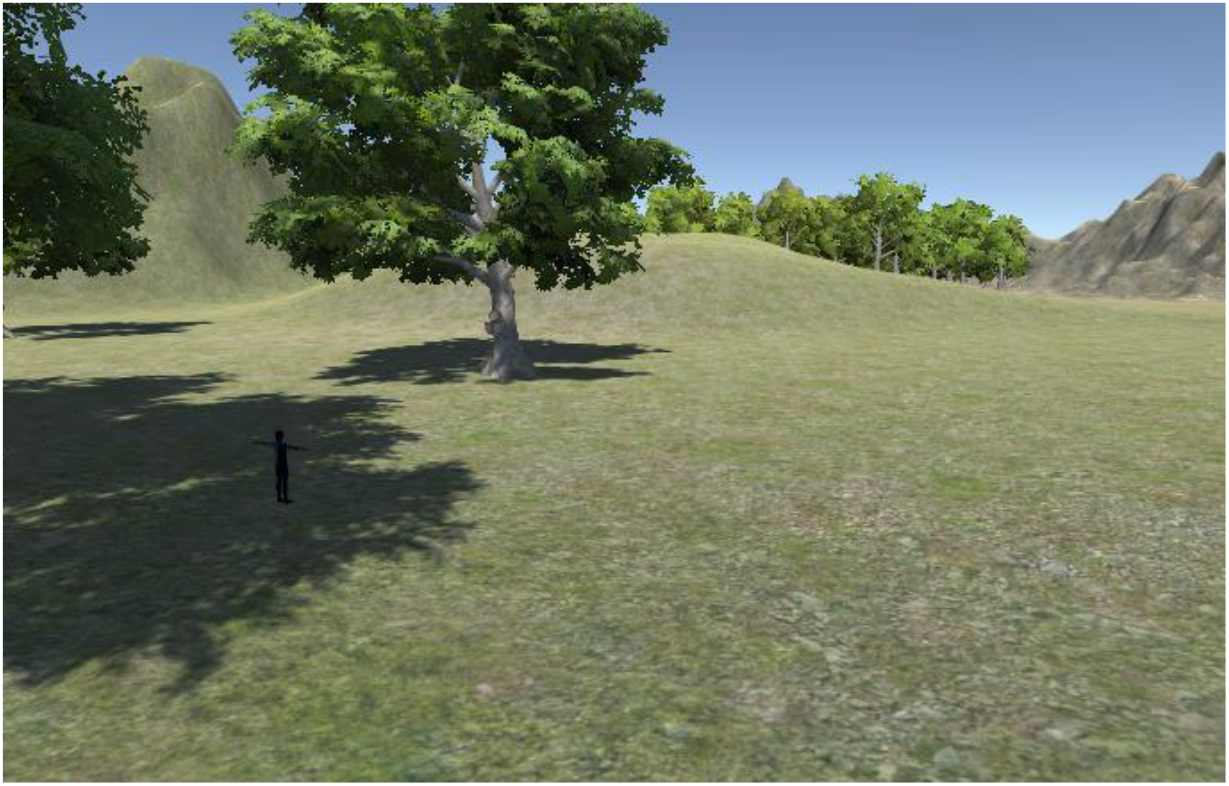
Example of a realistic, large-scale environment created using the terrain tool within Unity3D.

### 3.4. Physics

Unity’s physics system^14^ allows entities within the game to respond realistically to forces such as gravity and collisions with other objects. This is achieved by the simple addition of a Rigidbody component^15^ which can be used to alter aspects such as the object’s mass or the drag applied during movement. This is one way in which Unity can aid researchers to move away from abstract tasks and towards more immersive environments which better represent reality.

### 3.5. Navigation and pathfinding

Unity’s navigation system^16^ allows the user to create agents that can intelligently find their way around an environment using ‘navigation meshes’ that are automatically generated from the environment’s geometry. When moving to a particular destination, agents will move over the geometry of the environment, using the shortest or most efficient path and avoiding obstacles, including each other. VERSE makes extensive use of Unity’s navigation system in programming artificial intelligence agents (AIs) which act as sources of social information. Unity’s navigation system makes AI movement much easier for researchers to achieve than, for example, hard-wiring agents with a highly specific path. This also adds more realism, randomness and inter-individual variability to AI movement by allowing them to, within the constraints of the specified parameters, find their own path to their target location. VERSE can also extract the 3D location information of individuals at runtime, including the exact paths taken by AIs (see ‘Tracking values and logging data’, section 4.4.6, below).

### 3.6. Visual effects

Unity comes with a variety of visual effects that can add an extra dimension to the environment by increasing the level of realism. Examples that VERSE explicitly makes use of are lights^17^ and particle systems^18^. Lights are glowing effects that illuminate the surrounding objects, while particle systems are used to emit a number of small meshes to produce a visual effect (e.g. fire, smoke, water splashes) according to the properties inputted by the user.

### 3.7. Building an application with multiple scenes

Unity allows the user to build a game or application with multiple levels or scenes. For the purpose of human behavioural research, this can be a useful way of organising several replicates into a single application, so a participant can take part in all replicates one after the other without the researcher having to generate many different builds.

## 4. “Virtual Environments for Research into Social Evolution” (VERSE)

The remainder of this article will describe how I brought together these elements of Unity3D functionality to produce a custom tool for research. VERSE is a research tool, created using Unity3D (version 2017.4.23f1), designed to allow researchers to create their own realistic virtual environments for studying human social learning. VERSE is designed to be flexible, providing researchers with many different ‘building blocks’ that can be used to create tasks of different types, generate AIs with particular behaviours and allow environmental fluctuations. VERSE is also designed to give participants free movement over potentially large-scale environments, allowing experiments comparable to those involving wild animal populations or human hunter-gatherer communities to be produced. VERSE has been developed in a way that makes it widely available to all researchers, avoiding the use of costly equipment such as VR headsets and providing all the code necessary to build social learning environments without the need for game coding knowledge. While VERSE was specifically created with the study of human social learning in mind, it could feasibly be expanded to study other subjects, such as epidemiology or other aspects of human behaviour. VERSE can be accessed at the following figshare repository: https://figshare.com/s/c97c305736c9a3d1c8b9 (Easter, 2022) and requires Unity to open.

VERSE uses a combination of the standard Unity3D features, as described above, and a catalogue of specialised scripts coded specifically for the tool. Using VERSE will therefore require some basic knowledge of the Unity interface, but largely involves altering the properties of pre-created objects and so does not require any coding knowledge unless the researcher wishes to make substantial changes. What follows is an account of the novel features and their associated scripts coded specifically for VERSE (summarised in Figure 4.2). Script names are given in *italics*, VERSE-specific GameObjects are given in `courier font` and computer keys are highlighted in **bold**. A basic account of the function of each script is given in the main text. For researchers wishing to use VERSE, a detailed instruction manual / tutorial can also be found in the figshare repository.

**Figure 0.2.**
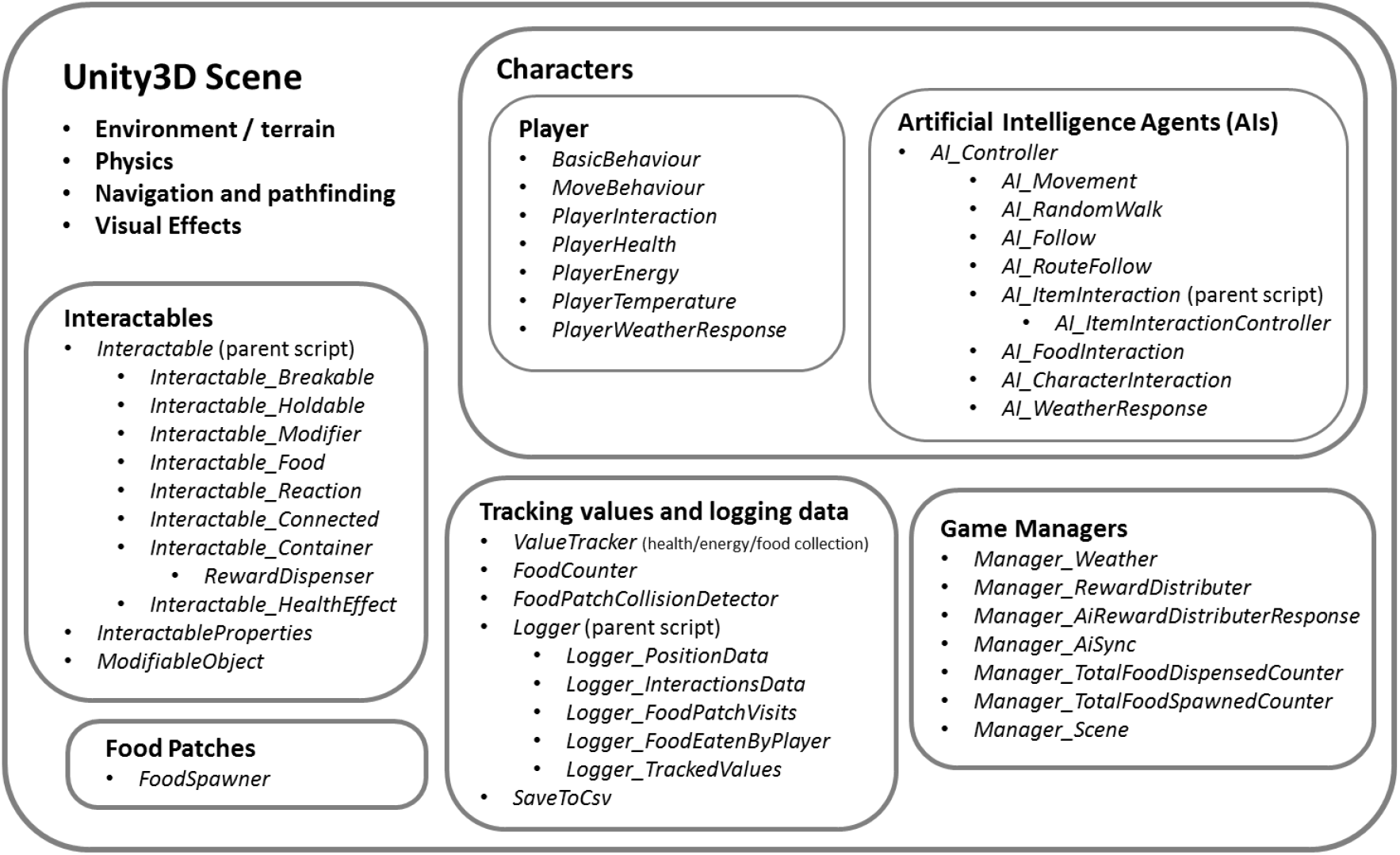
Schematic showing the main features of VERSE and the various scripts that control them. Script names are shown in italics and their descriptions and functioning can be found in the main body of the manuscript.

### 4.1. Characters

VERSE uses a third person character controller asset from the Unity Asset Store^19^, which is modified to produce two different types of character – the `player` (which is controlled by the participant) and Artificial Intelligence agents (`AIs`; controlled by the game code and programmable by the researcher). The `player` and `AIs` share the same general appearance and animations. Some of the animations were in-built features of the third person character controller asset^19^, including walking, running, jumping and idle animations. Other animations were created specifically for use in VERSE, including interacting and crouching animations. Both character types have a ‘collider’, which gives them a physical presence and stops them being able to walk through solid objects, and a ‘trigger’, which allows them to register when other objects are nearby (Table 4.1). The `player`, `AIs` and the scripts that can be applied to each of them are described in more detail below.

### 4.2. The Player

The `player` is the virtual human that a participant takes control of using the computer keys and is followed by a camera. The `player` is capable of various behaviours and can be affected by various elements of the game, according to the needs of the researcher. The scripts relating to `player` movement and behaviour are detailed below. Note that all computer keys described below are default options and can be changed via the Unity settings (e.g. to improve accessibility).

#### 1.1.1.1. BasicBehaviour and MoveBehaviour

Two scripts that work in conjunction to allow a participant to control the `player` using the computer keys. These scripts are slightly modified versions of those available in the third-person character controller^19^. Default `player` controls are given in Table 4.2. Movement, sprinting and jumping behaviours were all taken from the original code. Crouching behaviour is a VERSE addition – its only function is to serve as an additional behaviour that can be required to complete certain tasks (see ‘Interactables’, section 4.4.3, below). The options within the Unity interface allow the researcher to alter aspects of `player` movement, such as walk, run and sprint speed. An additional option allows the researcher to disable jumping, as this may be unnecessary or undesirable in certain studies.

**Table 0.2.** Default `player` controls according to the *BasicBehaviour and MoveBehaviour* scripts and Unity settings.

| Player control | Description | Computer key (default) |
| --- | --- | --- |
| Movement | Player moves along Unity’s (x,z) plane, which changes from walking to running depending on how long the key is held down. | Arrow keys<br>(←,↑,→,↓) |
| Sprint | Speeds up movement. | **Left Shift** |
| Crouch | Player crouches down. | **Left Ctrl** |
| Jump | Player jumps. Distance is further if walking or running at the time. | **Spacebar** |

#### 1.1.1.2. CameraControl

Script that is placed on a camera to allow camera rotation. The camera itself must be positioned correctly and connected to the `player` via Unity’s hierarchy system to allow player movement to be tracked by the camera (see instruction manual in the figshare repository for more information). The camera can be rotated left and right using the **A** and **D** keys, and rotated up and down using the **W** and **S** keys. The researcher can alter the vertical and horizontal rotation speeds within the Unity interface. In addition, depending on the task in question, the vertical rotation may be unnecessary or distracting in some studies and so can be switched off completely.

#### 1.1.1.3. PlayerInteraction

Script that allows the `player` to interact with specific ‘interactable’ objects (see ‘Interactables’ below). By default, this involves the `player` approaching the object and pressing the ‘Interact’ button (default: **?** key, modifiable in the Unity settings), however alternative requirements can be programmed for each individual interactable object, as discussed below.

#### 1.1.1.4. PlayerHealth

Stores a numeric value representing the `player`’s current health and displays this as a health bar onscreen, which updates during gameplay. This script contains functions to add or remove health from the `player` which can be called on by interactable objects. This script also logs when the `player`’s health reaches zero, which can be used to end the level or game prematurely (see *Manager_Scene*, section 4.4.7.7, below). Adding a health component to the `player` can be useful for assessing how humans learn behaviours based on their consequences. One way in which a `player`’s health can be affected is by collecting `Food interactables` (see *Interactable_Food* in the ‘Interactables’ section below, section 4.4.3.3.4) – poisonous `Food` that depletes health and/or nutritious `Food` that increases health by different amounts can be created and used to study human learning about food preferences. Thus, VERSE allows the possibility of creating tasks with clear evolutionary significance, comparable to foraging tasks used in animal studies.

#### 1.1.1.5. PlayerEnergy

Stores a numeric value representing the `player`’s current energy level and displays this as a bar onscreen, which updates during gameplay. Whenever the `player` moves (i.e. whenever the participant presses down any of the movement keys), the `player`’s energy depletes at a specified rate. When running (i.e. if the participant holds down the **Shift** key while moving), the energy depletion rate is multiplied by a specified ‘sprint multiplier’. The researcher can also specify how much energy should be used while jumping. Energy recovery can also be enabled – meaning that the `player`’s energy value will increase by a specified rate up to a specified maximum whenever the `player` is not moving, thus giving participants the option of ‘resting’ when their energy is low. The researcher can also specify whether energy depletion should be linked to player health. If player energy is linked to player health, once the `player`’s energy reaches zero, the `player`’s health starts to deplete at a specified rate. Adding an energy component is useful in tasks that require a participant to make energetically efficient choices, such as the route choice tasks, mazes or foraging tasks used in some animal studies (e.g. Laland and William, 1998). Allowing energy depletion to have an effect on health can further reinforce the choices that participants make.

#### 1.1.1.6. PlayerTemperature

Stores a numeric value representing the `player`’s current temperature, which is displayed as a temperature gage onscreen. The researcher can specify the values for the `player`’s ideal body temperature and for minimum and maximum possible temperatures, as well as the visual properties of the temperature bar. By default, the `player`’s body temperature is set to 37 and the minimum and maximum temperatures are set to 35 and 39, representing the temperatures that lead to hypothermia and hyperthermia in humans, but this can be changed according to the researcher’s requirements. As discussed below, the temperature of the `player` can be programmed to fluctuate in different ways according to different `weather conditions`, which can optionally be detrimental to health.

#### 1.1.1.7. PlayerWeatherResponse

Determines how the `player` reacts to changes in the ‘weather’ (see *Manager_Weather*, section 4.4.7.1, below). Weather can affect the `player` in the following ways: (i) raise/lower temperature at a specified rate; (ii) raise/lower health at a specified rate; or (iii) raise/lower temperature to a specified maximum/minimum (according to the *PlayerTemperature* script, see above), beyond which health begins to deplete at a specified rate. These effects only occur when the specified `weather condition` is currently active. Once the `weather condition` has deactivated, or if the `player` moves into a `weather shelter` (see *Manager_Weather* below), the `player`’s temperature gradually returns to the ideal body temperature. However, any health effects incurred by the `weather condition` remain even after the `weather condition` has been deactivated.

### 4.3. Interactables

This section refers to objects that can be interacted with in various ways by either the `player` or `AIs` with interaction capabilities enabled. To interact with an `interactable`, a character (the `player` or an `AI`) must approach it and meet a specified set of requirements for a successful interaction to occur. Each `interactable` is surrounded by a trigger^6^ which senses when an object moves through it (Table 4.1). The size of this trigger determines how close a character needs to be to the `interactable` in order to attempt an interaction (this will henceforth be referred to as the `interactable`’s ‘trigger area’). `Interactables` can also be ‘modified’ to affect their appearance and functionality (through the use of the *ModifiableObject* script; section 4.4.3.2), which provides VERSE with further flexibility for constructing novel tasks. Unless otherwise specified by the researcher, all `interactables` are coded to behave in the same way towards any character, whether it is a `player` or `AI`. This ensures that social information is realistic – i.e. a participant can observe an `AI` performing a particular set of behaviours and imitate these behaviours to achieve the same result. What follows are descriptions of all `interactable`-related scripts within VERSE.

#### 1.1.1.8. InteractableProperties

This script is automatically added to a GameObject when any one of the *Interactable* scripts (below) is attached, and holds information about the `interactable` in question, including its ‘item type’. This script also determines the fate of an `interactable` object once it has been destroyed (i.e. removed from the environment); which can occur in a number of circumstances, including if it is a `Breakable interactable` that has been broken or a `Holdable interactable` that has been deposited into a `Container` (see below). After an `interactable` object is destroyed, it can either be removed from the environment entirely or regenerated, either in its current position or in its original position, after a specified number of seconds. Having an `interactable` regenerate once it has been destroyed may be useful if, for example, the researcher wishes the participant to complete the same task multiple times in succession without creating multiple versions or replicates. This can be likened to situations in animal social learning research where the researcher routinely adds an object to the subject’s environment during the study period (e.g. Horner *et al*., 2010). Regeneration can also be synced with that other `interactables` (i.e. this object will only regenerate when a set of specified `interactables` are also regenerated), which is useful if a researcher wishes to add a set of `interactables` back into the environment only when they have all been removed.

#### 1.1.1.9. ModifiableObject

This script allows an `interactable` to be modified by the addition of a ‘`modification`’ (`interactables` with the ability to be modified in this way will henceforth be referred to as `Modifiable Objects`). `Modifications` can alter the `interactable`’s appearance and influence how it interacts with other `interactables` (via their ‘required items’, as discussed in the description of *Interactable,* below). For each `Modifiable Object`, the researcher specifies a list of ‘possible modifications’ that can be applied and how each of these should affect the `interactable`. Each `modification` can alter the appearance by the addition of extra object(s) or by switching the material (colour/texture) of the object. A researcher can create any number of novel `modifications` using a VERSE-specific menu in the Unity interface and have these `modifications` affect different `Modifiable Objects` in different ways. `Modifications` are stored as ScriptableObjects^10^, allowing them to be easily added to the *ModifiableObject* component of an object within the Unity interface. `Modifications` can also be added to a `Modifiable Object` during gameplay via interaction with an appropriate `Modifier interactable` (see *Interactable_Modifier*, section 4.4.3.3.3, below) making it possible to allow learning of cumulative behaviours within VERSE. An example of a `Modifiable Object` may include a ‘stick’ that can be modified using a ‘burning’ `modification` by the addition of a ‘flame’ at the end and a change in functionality.

#### 1.1.1.10. *Interactable* (parent script)

This script acts as a base (or ‘parent’) script for all `interactables` and dictates several properties and functions that are inherited by all the ‘child’ scripts listed below (and summarised in Table 4.3). For each `interactable` object, a particular set of conditions can be specified that must be fulfilled during an attempted interaction for that interaction to be successful. Firstly, the researcher can specify a set of ‘required keys’ that must be pressed when the `player` is within the `interactable`’s trigger area to attempt an interaction (by default, this is the ‘Interact’ key: **?**) and, secondly, a set of ‘required items’ that must be held by the `player` or `AI` during an attempted interaction for that interaction to be successful (e.g. a stick may be required to reach a high-up object). Each required item can either be a specific `interactable` object within the scene or can be a general item type, as identified by the ‘item type’ in an `interactable`’s *InteractableProperties* script (see above). For example, there are a group of ‘keys’ (all with the item type ‘key’) located in the environment which the `player` can pick up and use to attempt to open a ‘chest’. If the chest has a specific key object set as a required item, the `player` must pick up and use that specific key for the interaction to be successful. On the other hand, if the required item has a general item type of ‘key’, which matches the item type of all the keys in the environment, any one of those keys could be used to successfully open the chest. In addition, for each required item, the researcher can also specify a particular `modification` (see *ModifiableObject,* above) that must be applied for the condition to be met (e.g. a ‘stick’ must have the ‘burning’ `modification` to allow an interaction with a ‘flammable’ `interactable`). For `interactables` that are intended for `AI` use only, `player` interaction can be completely disabled.

The *Interactable* parent script contains functions that check whether all interaction conditions have been met and, if so, initiates a successful interaction. The response to a successful interaction depends on the type of `interactable` concerned, as explained below. What follows are the various types of `interactables` available in VERSE (summarised in Table 4.3). Each `interactable` type is a ‘child’ of the *Interactable* ‘parent’ script. When one of the following *Interactable* child scripts is added to an object, that object becomes an `interactable` of a specified type, ‘inheriting’ the properties and functions detailed above, while also containing their own individual functionality. Each type of `interactable` responds to a successful interaction in a different way and this response is dictated by its own properties, as specified by the researcher.

**Table 0.3:** Summary of the different types of `interactable` available in VERSE, how they respond to a successful interaction, and an example of use.

| Interactable type | Response to successful interaction | Optional responses | Example of use |
| --- | --- | --- | --- |
| Breakable | Object breaks and disappears. | Any number of 'held items' can be released from the object after it has been broken. | Breakable 'fruit' that releases 'seeds'. |
| Holdable | Object is picked up and held by the player or an AI. | n/a | A 'stick' that can be picked up by the player. |
| Modifier | Applies a specified 'modification' to a compatible Modifiable Object currently in the character's hand. | Modification can be either added to or removed from the Modifiable Object. | A 'fire' modifier that can change a held 'stick' into a flaming 'torch' by the addition of a 'burning' modification. |
| Food | Object is 'eaten' by the player, affecting their health. Disappears after interaction. | Different food objects can increase or decrease player health by different amounts. This can occur instantaneously or after a delay. | A 'poisonous seed' released by a breakable 'fruit' that lowers player health on interaction. |
| Reactable | A visual reaction is switched on. | Reaction can be a light or a release of 'particles'. Can switch on and off upon successive interactions, or alternatively switch on only once. | A 'fire' that releases sparks when interacted with. |
| Connected | A connected object is affected. Possible effect on connected object: GameObject is 'activated'; physics is enabled on object; interactable object is interacted with. | Connected object can be affected just once or multiple times. | Interaction with a 'tree' applies gravity to a separate 'fruit' object, causing it to fall to the ground. |
| Container | Holdable interactable currently in the character's hand is deposited into the Container, provided it is one of the Container's 'accepted objects'. | Can double up as a 'reward dispenser' that gives out reward(s) on deposition of a correct object. AI rewards can be disabled. | A 'vending machine' that gives out a 'food' reward on deposition of a 'coin'. |
| Health Effect | Adds or removes health when the player interacts. | Can be instantaneous or after a specified delay. | A poisonous 'flower' that lowers health that the player should learn to avoid. |

#### 1.1.1.10.1. *Interactable_Breakable* (child script, inherits from *Interactable*)

An `interactable` that breaks open after a successful interaction. The `interactable` is destroyed and replaced with a ‘broken parts’ GameObject that acts as a visual representation of the broken object. These broken parts can either remain indefinitely or disappear after a specified number of seconds. One element of social learning is the influence of the products of a demonstrator’s behaviour on observer behaviour (Heyes, 1994). Having the broken parts of a `Breakable` object remain may therefore be useful if the researcher wishes to explore this type of social learning (e.g. does finding the broken remains of a ‘fruit’ encourage participants to try breaking the same type of fruit open themselves?). However, if a large number of `Breakable` objects are to be present in the scene and the influence of their broken remains are not a focus of the research in question, their removal is recommended to keep computational costs down. A `Breakable interactable` can optionally contain a number of ‘held objects’ which are released once the `interactable` has been broken. Examples may include a ‘fruit’ that breaks open when interacted with to release a number of ‘seeds’, or a ‘barrel’ that breaks open to release a number of collectable ‘coins’.

#### 1.1.1.10.2. *Interactable_Holdable* (child script, inherits from *Interactable*)

An `interactable` that can be picked up and carried by a character (`player` or `AI`). Each character can hold a maximum of two items (one in each hand) at any given time and can therefore only successfully interact with a `Holdable interactable` if at least one of its hands is free. Objects held by the `player` can be released from the left or right hand by pressing the [ or ] key, respectively. `AIs` can be programmed to release `Holdable interactables` according to certain conditions (see *AI_ItemInteractionController*, section 4.4.5.6.1, below). `Holdable interactables` are especially useful because they can be specified as ‘required items’ for interacting with other `interactables` (see above). An example may be a ‘stick’ that can be picked up by the `player` or `AI` and used to interact with other objects.

#### 1.1.1.10.3. *Interactable_Modifier* (child script, inherits from *Interactable*)

An `interactable` that applies a specified `modification` to a `Modifiable Object` (see above). The researcher specifies the `modification` to apply and whether it should be added to or removed from the `Modifiable Object`. The `Modifier` can only modify compatible objects (i.e. a `Modifiable Object` with the same `modification` listed in its set of ‘possible modifications’). An example may be a ‘fire’ that, when interacted with by a character holding a compatible ‘stick’ object, modifies that stick by adding a ‘burning’ `modification`, which turns the stick into a flaming ‘torch’.

#### 1.1.1.10.4. *Interactable_Food* (child script, inherits from *Interactable*)

An `interactable` that can be ‘eaten’ (or collected) by the `player` or `AI`. Each `Food` item has a ‘nutrition value’, a ‘nutrition delay’ and a ‘food type’ specified by the researcher. Upon successful interaction, the `Food` object disappears and is considered eaten or collected by the character interacting with it. When a `player` with a *PlayerHealth* script attached eats a `Food` item, it can optionally affect the `player`’s health, adding the nutrition value to the `player`’s current health value after a delay equivalent to the nutrition delay. This effect on health can be either positive or adverse, allowing researchers to create foraging tasks comparable to studies on animals and hunter-gatherer communities, e.g. learning to avoid poisonous foods (Galef, 2009; Henrich and Henrich, 2010). By including a nutrition delay, one can assess how social information use is impacted by the time taken to process information about payoffs (e.g. in reality, the effects of eating a toxic or inedible food item would not necessarily be instantaneous) or by the unpredictability of rewards gained by using a particular behaviour (c.f. Caldwell and Eve, 2014). The nutrition value and food type variables are also used by `AIs` when making decisions on what `Food` items to eat (see *AI_FoodInteraction,* section 4.4.5.7, below), thus allowing specific social information to be conveyed to participants. An example of a `Food interactable` may be an edible ‘seed’ released from a `Breakable` ‘fruit’ `interactable`, which increases the `player`’s health when eaten. Food rewards are often given to animals completing tasks in social learning studies and the addition of `Food interactables` in VERSE offer a comparable way of rewarding human participants – although note that the *Interactable_Food* script could also be used to create collectable, non-food items (e.g. collectable coins). Additional functionality relating to `Food interactables`, such as logs that keep track of the number of `Food interactables` collected (see *FoodCounter,* section 4.4.6.2, below), are also available in VERSE to aid data collection and allow participants to see how well they are doing in a task.

#### 1.1.1.10.5. *Interactable_Reaction* (child script, inherits from *Interactable*)

An `interactable` that reacts to successful interactions with a visual display. This display can involve a light^17^ and/or particle system^18^. Successful interactions can either switch the display on and off (which may be more appropriate for a light or a looping particle system), or only switch it on (which may be more appropriate for a non-looping particle system that switches off itself after a certain amount of time). This type of `interactable` is entirely visual. It could be used to draw a participant’s attention to a particular area or simply to visually enrich the environment. The use of a light display could also be used to illuminate a dark area as part of a task. An example of this type of `interactable` may be a ‘campfire’ that is lit (by switching on a particle system designed to look like fire) after a character interacts with it while holding a burning stick.

#### 1.1.1.10.6. *Interactable_Connected* (child script, inherits from *Interactable*)

An `interactable` that affects a connected object upon successful interaction. The connected object can be affected in one of three ways: (i) the connected GameObject is ‘activated’, meaning it becomes visible if it isn’t already and all its components and behaviours are enabled^20^; (ii) the connected object’s Rigidbody component^7^ is enabled, allowing it to respond to physics forces including gravity; or (iii) the connected object, provided it is a VERSE `interactable`, is interacted with as though the `player` is interacting with it directly, but bypassing its usual conditions for interaction such as required keys or close proximity to the object. The researcher can additionally specify whether the effect on the connected object can occur multiple times or whether it should happen only once. Examples of where `Connected interactables` could be used include a ‘tree’ that, when interacted with, causes a connected ‘fruit’ object hidden in the canopy to drop by enabling gravity; or a ‘button’ that breaks open a connected `Breakable interactable` that would normally require a required item to be held upon interaction, thus offering an alternative, perhaps faster, way of completing the task.

#### 1.1.1.10.7. *Interactable_Container* (child script, inherits from *Interactable*)

An `interactable` into which `Holdable interactables` can be deposited. A list of ‘accepted objects’ can be specified. Only if a character is holding an accepted object while interacting with the `Container` will a successful interaction occur. A successful interaction causes the `Holdable interactable` currently in the character’s hand to be deposited into the `Container` and subsequently destroyed from the game scene entirely (the fate of the `Holdable interactable` after this is then determined by its *InteractableProperties,* as discussed in section 4.4.3.1, above). The researcher can optionally program a `Container` to give out a reward upon deposition of an accepted object. This requires the addition of an extra script, *RewardDispenser*, which contains a ‘DispenseRewards’ function that *Interactable_Container* can call on during a successful interaction:

#### 1.1.1.10.7.1. RewardDispenser

This script allows the researcher to specify a list of ‘possible rewards’ – where each possible reward consists of a GameObject and quantity – and a ‘reward location’. It contains the function ‘DispenseRewards’, which chooses one of the possible rewards at random and generates the chosen reward object in the specified quantity at the specified reward location. The ‘DispenseRewards’ function is called on by the *Interactable_Container* script during a successful interaction, provided the `Container` has been set to give out rewards. A reward may, for example, consist of a number of `Food interactables` that can then be collected by the `player`. In some cases, it may be useful to have ‘dynamic’ `reward dispensers`, where the possible rewards swap and change between ‘rounds’. For example, a researcher may give a participant the task of choosing between multiple `reward dispensers` over a number of rounds and may wish to investigate how social information is being used during this task. If the location of the best reward was always fixed to a particular `reward dispenser`, it would be difficult to determine whether the participant was using social information or simply learning which `reward dispenser` was more profitable. To disentangle these effects, the *Manager_RewardDistributer* can be used to randomly swap the possible rewards between a set of specified `reward dispensers` (see *Manager_RewardDistributer*, section 4.4.7.2, below).

#### 1.1.1.10.8. *Interactable_HealthEffect* (child script, inherits from *Interactable*)

An `interactable` that increases or reduces the `player`’s health by a specified amount after a specified delay upon successful interaction by the `player`. A `Health Effect interactable` could be used, for example, to investigate avoidance behaviour – e.g. an environment could feature ‘toxic’ plants or animals that reduce the `player`’s health when interacted with and that the participant must learn to avoid. Alternatively, `Health Effect interactables` that increase player health could be used to explore hypotheses concerning how humans use social information about medicinal plants, as in Henrich and Broesch’s (2011) field study on Fijian village populations.

**Interactables example**

To illustrate how interactable scripts work together to allow complex tasks to be created, consider the following example: A participant is located in an environment and given a task to open a small wooden barrel using only the tools available. Those tools include a stick and a campfire. The participant is required to learn the following behaviours in order: pick up the stick, use the campfire to light the stick, and use the burning stick to destroy the barrel (as illustrated in Figure 4.3). In VERSE, this is possible using the extensive ‘interactables’ system. The stick, campfire and box are all different types of `interactable`, each containing scripts derived from the *Interactable* parent script and each requiring particular conditions for a successful interaction to occur. The stick is a `Holdable interactable`, containing the *Interactable_Holdable* script, with ‘Interact’ and ‘Crouch’ as required keys and no required items. This means that, if the `player` approaches the stick and presses the ‘Crouch’ and ‘Interact’ keys, they will pick the stick up. The stick is also a `Modifiable Object` with one possible `modification`: ‘burning’, which alters the stick’s appearance by the addition of a ‘flame’ at the top. The campfire is a Modifier, containing the *Interactable_Modifier* script. It has ‘Interact’ as a required key and has no required items. It is capable of adding the ‘burning’ `modification` to any compatible `Modifiable Object`. In this case, this means that, if the `player` approaches the campfire and presses the ‘Interact’ key while holding the stick in hand, because the stick is a compatible `Modifiable Object`, the ‘burning’ `modification` will be added to it. The barrel is a `Breakable interactable`, containing the *Interactable_Breakable* script. It has ‘Interact’ as a required key and has the stick with the ‘burning’ `modification` as a required item. This means that the `player` must be holding the modified stick `interactable` for a successful interaction to take place and so must have completed the aforementioned behaviours in the correct order to break open the barrel. This example illustrates how different types of `interactable`, derived from the same parent script, can be used to assess cumulative learning through the use of the interaction conditions and object modifications available in VERSE.

**Figure 0.3.**
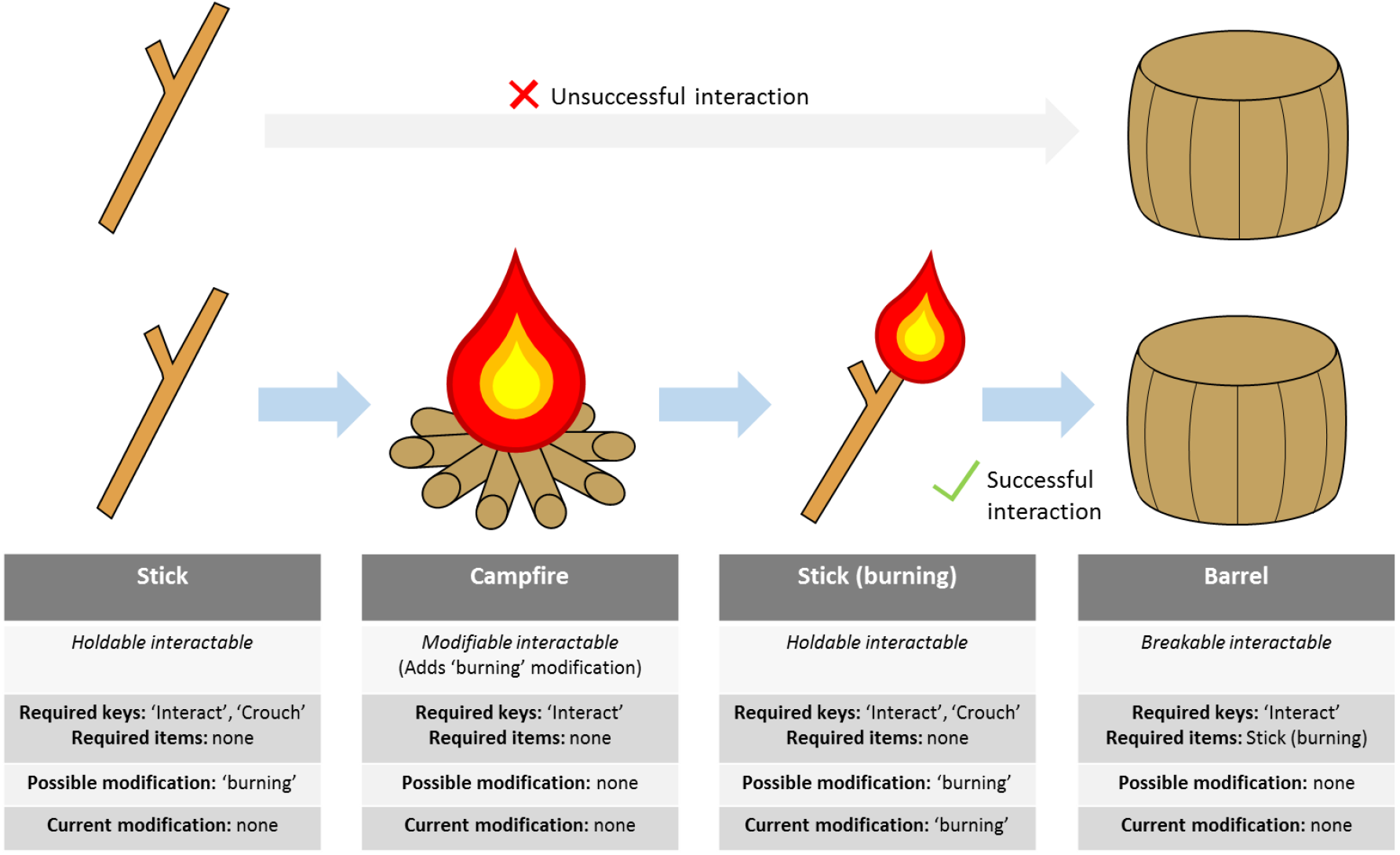
An example of how `interactables` work within VERSE. In this example, the participant is required to pick up the ‘stick’ (a `Holdable interactable`) and modify it with the ‘burning’ `modification` using the ‘campfire’ (a `Modifier interactable`) in order to break open the ‘barrel’ (a `Breakable interactable`) to receive a reward. Attempting to interact with the barrel while holding the unmodified stick results in an unsuccessful interaction. The participant therefore needs to use the correct behaviours in the correct order to complete the task.

### 4.4. Creating food patches

In VERSE, it is possible to create any number of `food patches` within the environment, which the `player` is required to find and exploit – thus allowing researchers to assess human foraging behaviour over large, complex envrionments. `Food patch` creation can be done simply by creating a number of `Food` objects and distributing manually around the environment. Alternatively, a researcher may wish to have `Food` items that are generated during gameplay and/or that regenerate after being ‘eaten’ (examples may include seasonal or temporal food supplies). This can be done using the *FoodSpawner* script.

#### 1.1.1.11. FoodSpawner

Spawns a specified number of a specified `Food` item at a specified ‘spawn location’. This occurs on a timer, with each spawn occurring after a specified time. Each individual `Food` item is spawned in a random position within a specified radius from the spawn location. The *FoodSpawner* can be programmed to spawn the `Food` items instantly at the beginning gameplay or wait until after the spawn time is reached before the first spawn occurs. It can also be programmed to reset its timer the instant it has spawned its last lot of `Food` items or, alternatively, wait until all the `Food` it previously spawned has become depleted before the timer is reset. The researcher can also limit the total number of `Food` items that can be spawned to a specified maximum. A number of *FoodSpawners*, each with their own specified `Food` items, can be added to the environment, or even to the exact same area to produce a patch of mixed `Food` items. It is worth noting that *FoodSpawners* can be programmed to spawn any object in this manner, not just `Food` items, and so can be repurposed for any task which requires objects to be generated on a timer during gameplay.

### 4.5. Artificial Intelligence agents (AIs)

In VERSE, social information about a task is conveyed in the form of Artificial Intelligence agents (`AIs`) with programmable behaviours. These `AIs` are designed to offer realistic sources of social information often not possible in lab-based or simple computer-based experiments by allowing participants to observe human figures moving around the environment and actively performing tasks, while also offering the researcher control over demonstrator behaviour and the social information being conveyed. A number of scripts are available in VERSE which control different aspects of `AI` behaviour, which can be added to each `AI` in different combinations depending on the behaviours that are appropriate for the given environment or task (e.g. for some environments, the researcher may only require movement-based behaviours, while others may require more complex behaviours).

#### 1.1.1.12. AI_Controller

This script is automatically added to a GameObject with any other `AI`-related script attached. Its purpose is to store a number of functions that offer overarching control over the `AI` and its behaviour and that can be called on by other `AI` scripts where appropriate. Functions include enabling and disabling `AI` scripts, pausing and unpausing `AI` behaviours and altering the parameters or properties of `AI` behaviours. The *AI_Controller* is especially important for ensuring that different `AI` behaviours do not interfere with one another.

#### 1.1.1.13. AI_Movement

A script containing functions used to instruct an `AI` to move to a specified destination at a particular walking speed, while updating character animation accordingly. The researcher can optionally specify whether an `AI` should move at an increased speed if they are moving further than a specified distance, which is a useful way of maintaining realistic movement speeds when `AI` behaviours span various distances. The functions in *AI_Movement* are called on by any `AI` scripts that require the `AI` to walk to a particular location.

#### 1.1.1.14. AI_RandomWalk

Programs an `AI` to move around according to a random walk that is controlled by several parameters inputted by the researcher. The researcher can also specify a set of ‘attractive areas’ – locations that are favoured in the random walk procedure over choosing a completely random point to walk to. For each attractive area specified by the researcher, the `AI` is given a ‘level of attraction’ which determines the probability that the `AI` will choose to visit that area. The researcher can also instruct the `AI` to favour less-visited attractive areas.

The random walk procedure is as follows. For each random walk cycle, the `AI` first waits for a random delay between a specified minimum and maximum. The `AI` then decides whether to visit an attractive area or walk to a random location. The probability of visiting an attractive area over choosing a random location is: 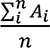 where *A_i_* is the level of attraction towards a particular attractive area, *i*, and *n* is the total number of attractive areas in the list. If the `AI` chooses to visit an attractive area, it then decides which area in its list it should visit. It iterates through all the attractive areas in its list and the probability of choosing attractive area *i* is: 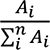. This formula ensures that an area is chosen and that areas further up in the list are not favoured over those at the bottom. If the researcher has instructed the `AI` to favour less-visited attractive areas, attractive area *i* will then be temporarily removed from the attractive areas list (thus ensuring it isn’t visited again) until all other attractive areas in the list have been visited. If the `AI` chooses not to visit an attractive area, or if no attractive areas are specified, the `AI` will instead walk a random distance between a specified minimum and maximum in a random direction from their its position. Movement to the chosen destination is executed via the *AI_Movement* script. Once the `AI` has reached its chosen destination, the random walk procedure is executed again, and this repeats continuously while ever the random walk script is enabled (i.e. unless it is disabled by the *AI_Controller* while another behaviour is being executed).

This random walk behaviour is designed to provide `AIs` with stochastic movement, while also allowing researchers to program `AIs` that provide social information about particular locations (e.g. foraging patches or other areas of interest). The ‘attractive areas’ feature could also be used for investigations into ‘local enhancement’, a social learning mechanism where a demonstrator’s behaviour attracts an observer to stimuli in a particular location (Heyes, 1994). In addition, the researcher can specify whether to allow `Food` detection during a random walk. When enabled, if an `AI` detects a compatible `Food` object while walking to its chosen location, it will abandon its current path and instead execute its `Food` interaction procedures (see *AI_FoodInteraction,* section 4.4.5.7, below). When disabled, an `AI` will ignore any `Food` items while on route to its chosen location. A researcher may use the first option if, for example, they are programming an `AI` that is designed to do a random search of the environment and stop off at any food sources that they pass by. Whereas the second option may be preferable if the researcher is programming an `AI` that is designed only to visit specific `food patches`, without becoming distracted by any other food sources it may pass.

#### 1.1.1.15. AI_Follow

Script that instructs an `AI` to walk to the location of a specified target object after a specified delay. Movement to this location is executed via the *AI_Movement* script. Once it has reached the location of the target object, the `AI` will repeat the behaviour. This behaviour can be used in conjunction with the random walk behaviour – the `AI` will switch between walking to the location of the target object and walking to a random location according to the respective time delays programmed into the two behaviours. One example of where this behaviour might be useful is if the researcher wishes to create a number of `AIs` that move as a group – one `AI` can be programmed with behaviours that determine the group’s chosen destination (e.g. via the *AI_RandomWalk* script) while the remaining `AIs` are instructed to follow the ‘leader’ using the *AI_Follow* script.

#### 1.1.1.16. AI_RouteFollow

Script that instructs an `AI` to follow a route through any number of specified waypoints, after an initial start delay randomly selected between a specified minimum and maximum number of seconds. This behaviour can be used in conjunction with the random walk behaviour – the random walk behaviour will be temporarily disabled via the *AI_Controller* script until the `AI` has followed the route to its end point. This behaviour is especially useful if a researcher wishes to investigate social information use during route-choice tasks, as has been investigated in both animals (Laland and Williams, 1998) and humans (Reader *et al*., 2008).

#### 1.1.1.17. *AI_ItemInteraction* (parent script)

Script containing functions to instruct an `AI` to interact with an `interactable`. If instructed to interact with a specified `interactable`, the `AI` will first check if the specified `interactable` is ‘available’. An `interactable` may become unavailable if (a) it is not present in the environment, e.g. if it has been destroyed, or (b) it is a `Holdable interactable` that is currently being held by the `player`, another `AI` or this `AI` itself. An `AI` will not attempt to walk to the location of or interact with an unavailable `interactable`. If the `interactable` is available, the `AI` will walk to the `interactable`’s location via the *AI_Movement* script, first disabling any random walk procedures via the *AI_Controller* script. The `AI` will then perform a second check to ensure the `interactable` has not become unavailable during the time the `AI` has taken to walk to it. If it is still available, the `AI` will attempt to interact with the `interactable`. As with the `player`, a successful interaction will only occur if the `AI` meets the required items condition for interacting. However, since the `AI` is computer-controlled, the required keys condition does not apply. The `AI` is assumed to be ‘pressing’ the correct keys and will play the appropriate animations associated with the required keys (e.g. playing the interaction animation if the ‘Interact’ key would be required by the `player`, or the crouching animation if the ‘Crouch’ key would be required). This provides a realistic source of social information, enabling participants to watch and imitate demonstrated behaviours to successfully complete a task.

#### 1.1.1.17.1. AI_ItemInteractionController (child script, inherits from AI_ItemInteraction)

This script gives the researcher control over what `interactables` an `AI` is able to interact with and how they interact with them, thus determining how social information about `interactables` is conveyed to participants. The researcher specifies a list of ‘potential interactables’ that the `AI` is able to interact with. For each `interactable` in the list, the researcher specifies whether the `AI` should interact with it only once or if multiple interactions are allowed. The basic procedure is as follows: An `AI` will interact with the `interactables` in its potential interactables list on a timer. After a random delay between a specified minimum and maximum number of seconds, the `AI` will choose an `interactable` (*i*) from its potential interactables list, then walk to its location and interact with it using functions inherited from the *AI_ItemInteraction* script. Once it has interacted with its chosen `interactable`, the timer will reset and the behaviour is repeated. If the researcher has specified that the `AI` should only interact with `interactable` *i* once, *i* is removed from the `AI`’s potential interactables list after the interaction occurs, regardless of whether said interaction was successful or not.

The researcher has a number of options concerning how an AI will choose between the interactables in its potential interactables list:

1. **Interact in order.** The `AI` will loop through the `interactables` in its potential interactables list in the order they have been inputted by the researcher within the Unity interface. This can be particularly useful if a researcher wishes to convey social information to a participant about cumulative behaviours or tasks that require multiple steps in a specific order to complete. The researcher can also specify whether the `AI` should continuously loop through its potential interactables list, returning to the first `interactable` once it has interacted with the last, or whether it should only iterate through the list once. In addition, the researcher can specify whether the `AI` should skip past any unavailable `interactables`, or whether it should wait for the next `interactable` in the list to become available again.
2. **Favour closest interactable.** The `AI` will calculate the distance between itself and all currently available `interactables` in its potential interactables list, choosing the one that is currently closest.
3. **Favour most recent interactable.** In this case, the `AI` will always choose the `interactable` with which it last interacted – provided that `interactable` is currently available, else it will choose its next `interactable` from its potential interactables list at random instead. This option may be useful, for example, if a researcher wishes to program an `AI` with a repertoire of possible behaviours, but have it preferentially repeat the same behaviour / option once it has been selected.
4. **Choose random interactable.** An `interactable` is selected at random from the currently available `interactables` in the `AI`’s potential interactables list.

An `AI` can also be programmed to add new `interactables` to its potential interactables list during gameplay if it comes in contact with them. An `AI` is considered as coming into contact with an `interactable` if it has entered the `interactable`’s trigger area. Having an `AI` add new `interactables` to its potential interactables list may be useful if a researcher wishes to add an element of randomness to an `AI`’s behaviour. Alternatively, a researcher may not wish the `AI` to interact with objects straight away at the start of the game and instead have the `AI` ‘learn’ about the `interactable` only after coming into contact with it, the likelihood of which can be controlled by the `AI` movement (e.g. by setting the location of the `interactable` as an attractive area, as described in *AI_RandomWalk*, above).

This script also instructs the `AI` how to treat any `Holdable interactables` it has picked up. An `AI` can carry a maximum of two `Holdable interactables` at a time (one in each hand). An `AI` will carry a `Holdable interactable` around indefinitely unless the researcher programs the `AI` to either deposit it into a `Container`, or drop it given a specified condition. An `AI` can be programmed to drop a `Holdable interactable` according to any of the following conditions: (a) after a random number of seconds between a specified minimum and maximum; (b) after a specified number of interactions with other `interactables` in their potential interactables list; or (c) after the `interactable` has been successfully ‘used’ a specified number of times – with a ‘use’ defined as a successful interaction with another `interactable`, for which this `Holdable interactable` and its modification(s) were a ‘required item’. Allowing an `AI` to pick up, use and drop `Holdable interactables` can be useful if a researcher wishes to create an environment where the `player` is required to observe how an `AI` uses a `Holdable interactable` and then attempt to replicate the demonstrated behaviour.

#### 1.1.1.18. AI_FoodInteraction

This script allows `AIs` to collect specific `Food` items in order to transmit social information about `Food` types to the `player` or allow `AIs` to collect rewards after completing tasks. Despite being a type of `interactable`, `Food` items are dealt with in a separate `AI` script than other `interactables` since different parameters and `AI` behaviours are required – and to allow the collection of `Food` items (which are likely to act as rewards for the successful completion of tasks) to be completely independent of interactions with other types of `interactable`.

An `AI` has a specified `Food` detection radius and can only react to `Food` items within this radius. After a specified delay, the `AI` will first pause the timer in the *AI_ItemInteractionController* script to avoid interactions being missed during `Food` consumption, then the `AI` will choose a `Food interactable` within its detection radius and interact with it. If the interaction is successful, the `Food` will be collected (or ‘eaten’) by the `AI`, as discussed in *Interactable_Food* (section 4.4.3.3.4) above. The actual interactions with `Food` items are dealt with via the *AI_ItemInteraction* script.

The researcher has a number of additional options concerning how an AI responds to Food items:

1. **High value preference.** The `AI` interacts with the `Food` items in its detection radius in order of their nutritional values – interacting with the highest value `Food` first, then the second highest, and so on.
2. **Avoid poisonous food.** The `AI` will ignore any `Food` item that has a negative nutritional value (i.e. a ‘poisonous’ `Food` that would deduct health from the `player` if the `player` was to eat it).
3. **Specific food preferences.** The `AI` will only respond to specified types of `Food` items within a ‘food preferences’ list specified by the researcher, and will ignore all other `Food` types. This can be a useful way of researching whether human food preferences are spread via social transmission (i.e. do people favour food types that they have witnessed another individual eating?), as has been investigated in various animal species.
4. **Leave food behind.** Instructs an `AI` not to eat all the `Food` items in the area. An `AI` will only respond to `Food` if there are more than a specified number of `Food` items in its detection radius. This option is useful in situations where an informed `AI` is likely to visit a `food patch` before the uninformed `player`, to ensure there is still `Food` available when the `player` arrives.

#### 1.1.1.19. AI_CharacterInteraction

Script that instructs the `AI` to respond in certain ways towards other `AIs` and the `player`. `AIs` can respond to other characters in three ways: an aggressive display (stand up straight and punch outwards); a submissive display (hunch over, shield face and turn away); and a positive display (waving). When the `AI` comes within a certain distance of a character (i.e. within its trigger area), they turn to the character and, after a specified delay, perform one of the displays above. The type of display towards the `player` is explicitly chosen by the researcher, meaning a particular `AI` will always respond in the same way to the `player`. The `AI` also has a specified ‘aggression level’, which determines how it acts towards other `AIs`. If two interacting `AIs` both have an aggression level of zero, each with interact with a positive display. If both have aggression levels above zero and there is a difference in their aggression levels, the `AI` with the highest aggression level will use an aggressive display and the one with the lowest aggression level will use a submissive display. If both have an aggression level of above zero, but their aggression levels are equal, they will both use an aggressive display. If one of the `AIs` does not possess the *AI_CharacterInteraction* script, no interaction will occur. Alternatively, the researcher can specify a ‘specific response’ towards a particular `AI`, which will override any response based on aggression levels.

Previous research has that both human and non-human individuals are more likely to copy demonstrators with particular characteristics, such as dominant or familiar individuals (e.g. Henrich and Henrich, 2010; Horner *et al*., 2010; Kendal *et al*., 2015; Corriveau and Harris, 2009; Swaney *et al*., 2001; Guillette *et al*., 2016). Creating `AIs` within VERSE that interact with other characters using particular displays can enable participants to distinguish between `AIs` of different characteristics or personalities and investigate what influence this has on social information use. In addition, `AI` aggression levels and the resulting behaviours towards one another can also be used to generate a dominance hierarchy, which have been experimentally demonstrated as having an influence on social learning in animals (e.g. Kendal *et al.,* 2015).

#### 1.1.1.20. AI_WeatherResponse

Determines how an `AI` responds to different `weather conditions` (see *Manager_Weather*, section 4.4.7.1, below). The researcher specifies which `weather condition(s)` the `AI` will respond to and a list of ‘known weather shelters’. When one of these `weather conditions` is activated, the `AI` will, after a specified delay, run to one of the `weather shelters` in its list and stay in that shelter until the `weather condition` is deactivated. All other behaviours are disabled via the *AI_Controller* while an `AI` is shielding itself from the weather. The researcher can also specify whether the `AI` should favour the closest of its known shelters. This behaviour allows researchers to supply participants with social information about appropriate responses to unfavourable environmental conditions within environments where these `weather conditions` have been programmed.

### 4.6. Tracking values and logging data

Several scripts are available in VERSE for collecting and exporting various types of data that correspond to different types of task used in social learning research.

#### 1.1.1.21. ValueTracker

Tracks a specified numeric value during gameplay, updating continuously and additionally recording the value as a percentage of a ‘maximum value’, which is calculated differently depending on the type of *ValueTracker. ValueTrackers* are useful for continuously monitoring important information that the researcher may, for example, wish to display to the participant during gameplay. There are currently three types of *ValueTracker*:

1. **Player energy tracker.** Requires the *PlayerEnergy* script to be attached to the `player`. Tracks the `player`’s current energy as a percentage of the starting energy value. This can be used to monitor how energy-efficient a participant’s choices are – for example, in a route choice task where a number of routes of different length are available.
2. **Player health tracker.** Requires the *PlayerHealth* script to be attached to the `player`. Tracks the `player`’s current health as a percentage of the maximum health. This can be used to monitor how adaptive a participant’s behavioural choices are – for example, in a foraging task where a number of possible `Food` types, including potentially poisonous items, are available.
3. **Food collection tracker.** Tracks the number of `Food` items obtained by the `player` as a percentage of the total number of available `Food` items in the environment. This type of *ValueTracker* requires an additional *FoodCounter* script (see below) and updates its tracked value according to the current `player` food count. In environments where `Food` is generated during gameplay, calculating the total number of available `Food` items will require the *Manager_TotalFoodDispensedCounter* and/or *Manager_TotalFoodSpawnedCounter* scripts, depending on the way in which `Food` items are added to the environment (see sections 4.4.7.5 and 4.4.7.6, below). The food collection tracker is a useful way of assessing how ‘well’ a participant is doing in terms of food / rewards collected.

#### 1.1.1.22. FoodCounter

Counts the number of `Food` items collected by the `player` within the current environment. The researcher can specify whether this count should include the nutritional information of `Food` items or whether it should be based solely on the number of items collected.

#### 1.1.1.23. FoodPatchCollisionDetector

Detects when a character visits a `food patch` (or any other specified area). This script must be added to a GameObject that has a trigger^6^ around it (Table 4.1).

#### 1.1.1.24. *Logger* (parent script)

Creates a log of a specific dataset in preparation for export from the application.

#### 1.1.1.24.1. *Logger_PositionData* (child script, inherits from *Logger*)

Records the position (x, y, z coordinates) of a specified set of monitored objects. The position of each object and the current time in seconds is logged throughout gameplay at a specified time interval. A separate dataset is created for each monitored object in preparation for export. This can be used to assess how a participant uses social information about particular locations or routes. (Note: In Unity, y coordinates corresponds to the up/down direction, while x and z are the left/right and forward/backward directions, which may differ from statistical programs).

#### 1.1.1.24.2. *Logger_InteractionsData* (child script, inherits from *Logger*)

Records any interactions that a specified set of monitored characters (i.e. `player` or `AIs`) make with a specified set of monitored `interactables`. Each time a character successfully interacts with an `interactable`, the current time, the character name and the `interactable` name are recorded. One dataset is produced containing all the interactions made by all the monitored characters towards any monitored `interactable`, in the order at which they occurred. This can be used to assess how a participant uses social information about particular `interactables`.

#### 1.1.1.24.3. *Logger_FoodPatchVisits* (child script, inherits from *Logger*)

Records each time a character from a set of specified monitored characters visits one of a set of specified `food patches`. Requires a *FoodPatchCollisionDetector* script (section 4.4.6.3) to be added to each monitored `food patch` to register visits. A single dataset is produced showing all the visits made by any of the monitored characters to any of the monitored `food patches` and the times at which the visits occurred. This can be used to assess how a participant uses social information about the locations of `food patches.` (Note: This *Logger* can be repurposed to log visits to any area according to the researcher’s needs).

#### 1.1.1.24.4. *Logger_FoodEatenByPlayer* (child script, inherits from *Logger*)

Records all the `Food` items ‘eaten’ by the `player` during the course of the game. The name of the `Food` object, its nutritional value and the time at which it was eaten are all recorded. In addition, a cumulative nutritional value across all consumed `Food` items is recorded. If `AIs` are programmed to prefer certain food types, this *Logger* can be a useful way of assessing whether participant choices have been influenced by `AI` preferences.

#### 1.1.1.24.5. *Logger_TrackedValues* (child script, inherits from *Logger*)

Records the final tracked value, maximum possible value and percentage value of any *ValueTrackers* (see *ValueTracker* above) at the end of the current scene.

#### 1.1.1.25. SaveToCsv

Takes the dataset from all *Loggers* in the current scene and exports each as a .csv file at the end of gameplay. All datasets are exported to the game’s directory with descriptive file names to make data easy to recognise and sort. The researcher can also give participants a ‘reference number’ and have this number automatically added to all file names to allow data from the same participant to be linked together. VERSE comes with a dedicated scene where participants can input their reference number, which can be added to the start of the build – the *SaveToCsv* script will then save a record of this inputted number and add it to the filename of every outputted dataset.

### 4.7. Game managers

Several game managers are available in VERSE which add additional features to the environment as a whole and/or allow a greater level of control over some of the functions already discussed.

#### 1.1.1.26. Manager_Weather

Controls `weather conditions` within a VERSE environment. Each `weather condition` is saved as a ScriptableObject^10^ containing the following properties: *visual effect, duration, time interval* and *initial delay*. The *Manager_Weather* takes any number of `weather conditions` and enables and disables them on a loop according to their respective properties. Each `weather condition` has its own timer within the *Manager_Weather*. The `weather condition` is enabled after *time interval*, except at the beginning of the scene, when it is enabled after a delay equivalent to: *initial delay* + *time interval.* Once enabled, the `weather condition` is disabled again after *duration*, after which the timer restarts. Enabling a `weather condition` involves ‘switching on’ the *visual effect* associated with it. Visual effects for `weather conditions` are created using particle systems^18^ that are placed directly in front of the camera so they are continuously visible to the participant while ever the `weather condition` is enabled, without the high computational demands of creating a visual effect over the entire environment. The effects of the weather on the `player` and `AIs` can be altered through the *PlayerWeatherResponse* (section 4.4.2.7) and *AI_WeatherResponse* (section 4.4.5.9) scripts as previously discussed.

Any number of `weather shelters` can be added to an environment. These are areas or objects which protect the `player` from all `weather conditions` and which `AIs` can be programmed to run for when a particular `weather condition` is enabled. Each `weather shelters` is surrounded by a trigger^6^ (Table 4.1) which senses when a character enters it. Adding `weather conditions` to a social learning environment can be used both as a task in itself to establish how participants use social information to learn about adverse environments or to increase the difficulty of another, unrelated task through environmental complexity and uncertainty. This function aids VERSE in generating realistic environmental features only currently possible in field studies, while providing researchers with experimental control only possible in lab studies.

#### 1.1.1.27. Manager_RewardDistributer

Gives additional control over a set of specified `reward dispensers` (see *RewardDispenser*, section 4.4.3.3.7.1, above), allowing their rewards to be swapped and changed over a number of ‘rounds’. The researcher specifies a set of `reward dispensers` and, for each one, a set of ‘possible rewards’. The researcher also specifies the total number of rounds and can optionally have this displayed as text onscreen for the participant to see during gameplay. At the beginning of each round, the *Manager_RewardDistributer* randomly selects a reward for each `reward dispenser` from its set of possible rewards. This reward will override any possible rewards already set in the *RewardDispenser* script of the specified `reward dispenser`. The researcher can optionally specify that equal rewards should be distributed to all `reward dispensers`, in which case the *Manager_RewardDistributer* will randomly select a single reward from all sets of possible rewards and distribute this reward to all `reward dispensers`.

The *Manager_RewardDistributer* can also be programmed to swap the set of possible rewards between `reward dispensers` at a specified probability (where a probability of 1 means a definite swap) at the beginning of each round. The specific reward for each `reward dispenser` is then chosen as described above from the newly distributed sets of possible rewards. When swapping rewards between more than two `reward dispensers`, the set of possible rewards from each `reward dispenser` is distributed to one of the alternative `reward dispensers` at random. Programming a situation where rewards are swapped between rounds can allow a researcher to create a task where a participant is required to locate the `reward dispenser` with the highest reward and where the location of the highest reward is randomised, thus preventing the participant from simply learning a single location.

The researcher is also required to specify how the *Manager_RewardDistributer* should define the beginning of a new round. A new round can begin on a timer after a specified number of seconds or can be synced with the regeneration of a specified `interactable` (see *InteractableProperties*, section 4.4.3.1, above). An example of a situation where the second option would be favourable is a task where a `Holdable` ‘token’ must be deposited into one of two `Containers` to receive a reward, and where the token has been set to regenerate in its original position after being placed into a container. In this case, it may be favourable to sync the redistribution of rewards between the two containers with the regeneration of this token, therefore ensuring that the beginning of a new round, and so any change in the rewards dispensed by the containers, only occur after the participant has made their choice.

Once the total number of rounds have been completed, all rewards are removed from the set of `reward dispensers` and no new rounds will be initiated. In addition, the *Manager_RewardDistributer* can optionally be instructed to remove a number of objects from the scene once all rounds are completed. This can be useful in a situation where the beginning of each round is defined by the regeneration of an `interactable` object. When all rounds are completed, said `interactable` may become redundant and having it disappear from the environment can make it clear to the participant that the task is complete.

#### 1.1.1.28. Manager_AiRewardDistributerResponse

Controls how `AIs` respond to the reward distributions in the *Manager_RewardDistributor* script. A set of specified `AIs` are instructed to choose between a set of specified reward dispensers after a specified number of rounds. The researcher has several options influencing how `AI` choice is distributed between the specified reward dispensers:

1. **Frequency-distributed.** The researcher specifies the frequency of `AIs` that should visit each `reward dispenser`, and `AIs` are distributed randomly according to this frequency at the beginning of each round. The specified frequencies can either be fixed to a particular `reward dispenser` across all rounds (e.g. dispenser A is always visited by two `AIs` and dispenser B by three `AIs`) or can be set to swap randomly between rounds. Note that a 1:1 frequency can be used if a researcher wishes to investigate biases towards particular demonstrators while controlling for the frequency of demonstrators using choosing each option.
2. **Payoff-based.** Each `AI` is given a ‘tracking ability’ that determines how well it is able to detect the `reward dispenser` with the highest payoff. At the beginning of the round, each `AI`’s probability of choosing a `reward dispenser` with a higher payoff over one with a lower payoff is equal to its tracking ability.
3. **Frequency-distributed and payoff-based.** The researcher specifies the frequency of `AIs` that should visit each `reward dispenser` and `AIs` are distributed according to this frequency at the beginning of each round. The tracking ability of each `AI` acts as a weight that determines the likelihood that it will be chosen over other `AIs` to visit a higher-payoff `reward dispenser` within the specified frequency distribution. The frequencies assigned to each `reward dispenser` can be either fixed across rounds, set to swap randomly, or distributed according to payoff (so that the highest-payoff `interactable` is always visited by the most `AIs`).
4. **Assigned at random.** Each `AI` randomly chooses one of the specified reward dispensers at the beginning of each round.

For each `AI`, the chosen `reward dispenser` is added to the list of potential interactables within its *AI_ItemInteractionController* script (see section 4.4.5.6.1, above), allowing the `AI` to interact with the `reward dispenser` according to the rules and functions of this script. Increasing the number of rounds that elapse before choices are redistributed among `AIs` can be used to represent a delay in demonstrators responding to changes in their environment, thus adding a level of uncertainty about the social information being provided.

Overall, this manager allows the researcher to control how social information about reward dispensers is conveyed to the participant in a way that is specifically designed for investigating human social learning strategies. Frequency-, success- and model-based social learning biases have been demonstrated in many species (e.g. Pike and Laland, 2010; Kendal *et al*., 2009; Pike *et al*., 2010; Seppänen *et al*., 2011; Horner *et al*., 2010; Kendal *et al*., 2015), including humans (e.g. Morgan *et al*., 2012; Efferson *et al*., 2008; Mesoudi, 2008; Mesoudi and O’Brien, 2008; Henrich and Henrich, 2010; Miu *et al.,* 2018), but are generally difficult to assess in humans in a realistic environment, using ecologically relevant tasks. VERSE makes it possible to test such hypotheses within complex and naturalistic, yet highly controlled, environments.

#### 1.1.1.29. Manager_AiSync

Synchronises the item interaction behaviours within the *AI_ItemInteractionController* scripts of a specified set of `AIs`. For each `AI`, the researcher specifies a particular ‘sync interactable’ from the potential interactables list in its *AI_ItemInteractionController* script. When an `AI` interacts with this specified `interactable`, any further item interactions normally dictated by the *AI_ItemInteractionController* script are paused (via the *AI_*Controller) until all `AIs` have interacted with their own sync `interactables`. When all `AIs` have reached their sync `interactable`, item interaction is enabled again. Syncing `AI` item interaction behaviour is particularly helpful in cases where multiple `AIs` are required to display multiple options to the participant at the same time and/or where differences in the timing of `AI` choices may cause confusion or influence the choice of the participant.

#### 1.1.1.30. Manager_TotalFoodDispensedCounter

Calculates the maximum possible reward value that a `player` could collect from `reward dispensers` in the environment, which is used to update the maximum value in any food collection *ValueTrackers* (see *ValueTracker,* section 4.4.6.1, above) and to assess how ‘well’ a participant has completed a task involving `reward dispensers`. The researcher can specify which `reward dispensers` to monitor or, alternatively, specify that all `reward dispensers` in the environment should be included. The researcher can also specify whether `AI` rewards should be included in the total. A researcher may wish to include `AI` rewards in the total if they consider rewards dispensed to `AIs` as ‘extra opportunities’ for gaining rewards (e.g. if investigating ‘scrounging’ behaviour, which has been most notably demonstrated in pigeons; Giraldeau and Lefebvre, 1987). However, if a researcher wishes to base this total only on successful completion of tasks by the `player`, they may wish to avoid including `AI` rewards in the count.

The maximum number of rewards from `reward dispensers` is calculated as follows: `reward dispensers` are defined as ‘managed’ or ‘unmanaged’ depending on whether their rewards are managed by a *Manager_RewardDistributer* script (section 4.4.7.2). For unmanaged `reward dispensers`, the total number of possible `Food` rewards equate to the actual number of rewards dispensed during gameplay. Therefore, each time an unmanaged `reward dispenser` dispenses a reward into the environment, the reward value is added to the running total. For managed `reward dispensers`, rewards are distributed between a number of `reward dispensers` across a number of rounds (see *Manager_RewardDistributer*, section 4.4.7.2) and the participant is expected to choose only one `reward dispenser` per round. Therefore, for each round, only the maximum reward distributed amongst the managed `reward dispensers` (i.e. the maximum possible reward a `player` would gain if they chose the ‘best’ `reward dispenser` in the set) is added to the running total. If a researcher specifies that `AI` rewards should be included, any additional rewards dispensed to `AIs` from managed dispensers are viewed as opportunities to scrounge and so are added to the total as well. When calculating the total possible value from dispensed rewards, *Manager_TotalFoodDispensedCounter* can either count `Food` rewards by their quantity alone or can include each `Food` item’s nutritional value. This is determined by the options inputted into the *FoodCounter* script (see section 4.4.6.2, above).

#### 1.1.1.31. Manager_TotalFoodSpawnedCounter

Calculates the number of `Food` rewards spawned from any *FoodSpawners* in the environment (see *FoodSpawner,* section 4.4.4.1, above) during gameplay, which is used to update the maximum value in any food collection *ValueTrackers* (see *ValueTracker,* section 4.4.6.1, above) and to assess how ‘well’ a participant has completed a foraging task. `Food` rewards can either be counted based on their quantity alone or based on each `Food` item’s nutritional value. This is determined by the options inputted into the *FoodCounter* script (see section 4.4.6.2, above).

#### 1.1.1.32. Manager_Scene

Automates the process of moving to the next scene and/or ending gameplay so that the researcher does not need to do this manually. The application is instructed to end the current scene according to one or more of the following conditions: (i) after a specified amount of time has elapsed since the beginning of the scene; (ii) a specified delay after a *Manager_RewardDistributer* (see section 4.4.7.2, above) has completed all its rounds; (iii) if the `player`’s health reaches zero. Once a scene has ended, all functionality is paused pending the beginning of the next scene. If the current scene is the last in the build, the *Manager_Scene* displays an ‘end game canvas’, a 2D canvas^8^ (Table 4.1) that lets the participant know they have completed all tasks in the application. If the current scene is not the last in the build, the *Manager_Scene* can either move automatically on to the next scene, or optionally display an ‘end scene canvas’ which contains a button that allows the participant to decide when to move to the next scene. An end scene canvas may be preferred if the researcher wishes to allow participants a short break between scenes. In addition, one or more tracked values (see *ValueTracker*, section 4.4.6.1, above) can be optionally displayed on the end scene canvas and/or end game canvas, either as an absolute value or as a percentage of its maximum, to give the participant an idea of their performance during tasks.

## 5. Worked example

What follows is an example of a social learning environment created in VERSE (shown in Figure 4.4), which demonstrates some of the features described above. In this example, a single participant is placed into a small forest environment and given the task to find and break open fruits to collect seeds. The fruits are located in three trees within the environment and require a stick to reach and dislodge them. Hidden in the environment is a pile of sticks the participant can use. Two `AIs` provide a source of social information – both know the location of the stick pile and how to access fruit and break them open to get to the seeds, but each `AI` chooses to gather their fruit from a different tree. Data on the location of the `player` and `AIs` and all the `interactables` each character interacted with throughout gameplay are collected using the *Logger_PositionData* and *Logger_InteractionsData* scripts. This example environment could be used to investigate how social information is used when learning to use tools in a novel, realistic environment – and offers some direct comparisons to research on tool-use and foraging in animal communities (e.g. tool-use in chimpanzees; Biro *et al.,* 2003; Musgrave *et al.,* 2016).

**Figure 0.4.**
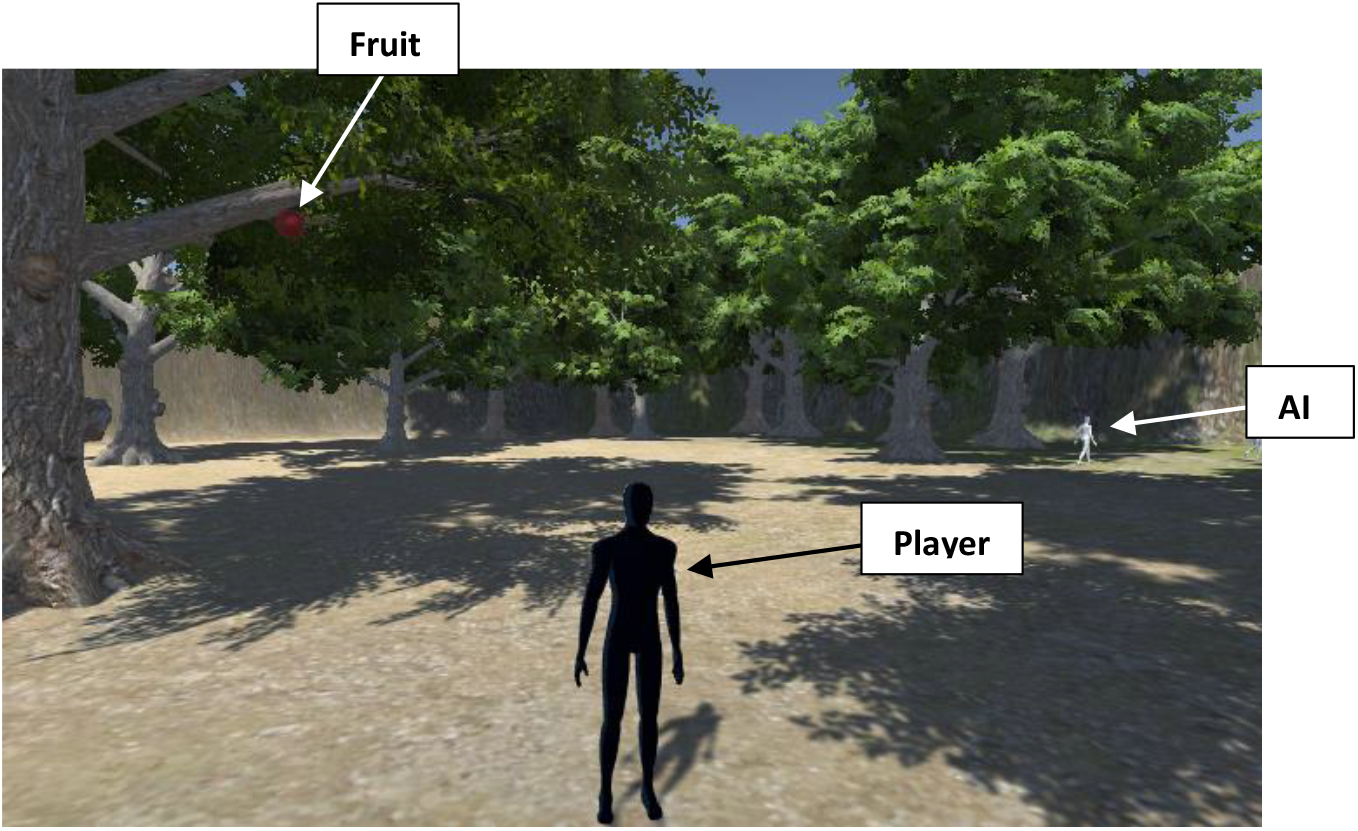
Example of a VERSE environment where the `player` (black figure, controlled by the participant) is required to obtain seeds by accessing and breaking open fruit hidden in trees. Computer-controlled `AIs` (white figures) provide social information about how to complete the task.

To achieve this, the environment is first created in VERSE using Unity’s terrain tool. A `player` is added to the environment that has the following three scripts attached: *BasicBehaviour, MoveBehaviour* and *PlayerInteraction*, which allow the participant to move the `player` and interact with `interactable` objects within the environment. A camera is added to the `player`, which follows the `player` around during gameplay. The *CameraControl* script is attached to the camera to allow camera rotation.

Three types of `interactable` are added to the environment: sticks, trees and fruit (Figure 4.5). Each `interactable` contains the *Interactable_Properties* script and one of the *Interactable* child scripts, as described below. A number of sticks are hidden in the environment. Three trees are placed in the environment, each with a fruit hidden in their branches. The fruit `interactables` have their ‘Rigidbody’ component disabled so they do not respond to gravity and so remain in the branches of the tree and out of reach of the `player` or `AIs`.

Each stick is a `Holdable interactable`, containing the *Interactable_Holdable* script (Figure 4.5A). One required key (‘Interact’) and no required items are specified, meaning the `player` simply needs to approach the stick and press the ‘Interact’ key (default: **?**) to interact with it. A successful interaction results in the `player` picking up and carrying the stick. The stick also has an ‘item type’ of ‘Stick’ specified in its *Interactable_Properties* component.

Each tree is a `Connected interactable`, containing the *Interactable_Connected* script (Figure 4.5B). It has one required key (‘Interact’) and one required item (any `interactable` with the general item type ‘Stick’). This means that, in order to interact with the tree, the `player` must approach the tree with a stick in hand (any stick with the correct item type, since a general item type has been specified rather than a specific object) and press the ‘Interact’ key for a successful interaction to take place. If the `player` attempts to interact with the tree without the required item in hand, the interaction will be unsuccessful. The *Interactable_Connected* script is programmed so that a successful interaction will enable the Rigidbody component of a connected fruit object, allowing the fruit to respond to forces such as gravity. This means that, when the `player` successfully interacts with the tree using the stick, the fruit will drop out of the tree, thus allowing the `player` to access it and break it open.

Each fruit is a `Breakable interactable`, containing the *Interactable_Breakable* script (Figure 4.5C). It has one required key (‘Interact’) and no required items. It is also set to release six ‘seeds’ when broken. This means that the `player` needs to approach the fruit and press the ‘Interact’ key to break open the fruit and gain access to the seeds. The seeds themselves are examples of `Food interactables`, containing the *Interactable_Food* script. Seeds have no required keys or required items, meaning the `player` simply needs to walk into them to collect them. To gain access to the seeds, the `player` must therefore learn to perform the correct series of behaviours, first picking up the stick and then using the stick to interact with the tree so that the fruit drops to the floor, then breaking open the fruit to access the seeds. The fruit also has an additional property specified in its *Interactable_Properties* script – it is set to regenerate in its original position after it is destroyed (i.e. when it is broken open, it will reappear back in the tree again). This gives the impression that the fruit ‘regrows’ after being hit out of the tree with the stick and gives both the `player` and `AIs` multiple opportunities to access fruit from the same tree.

**Figure 0.5.**
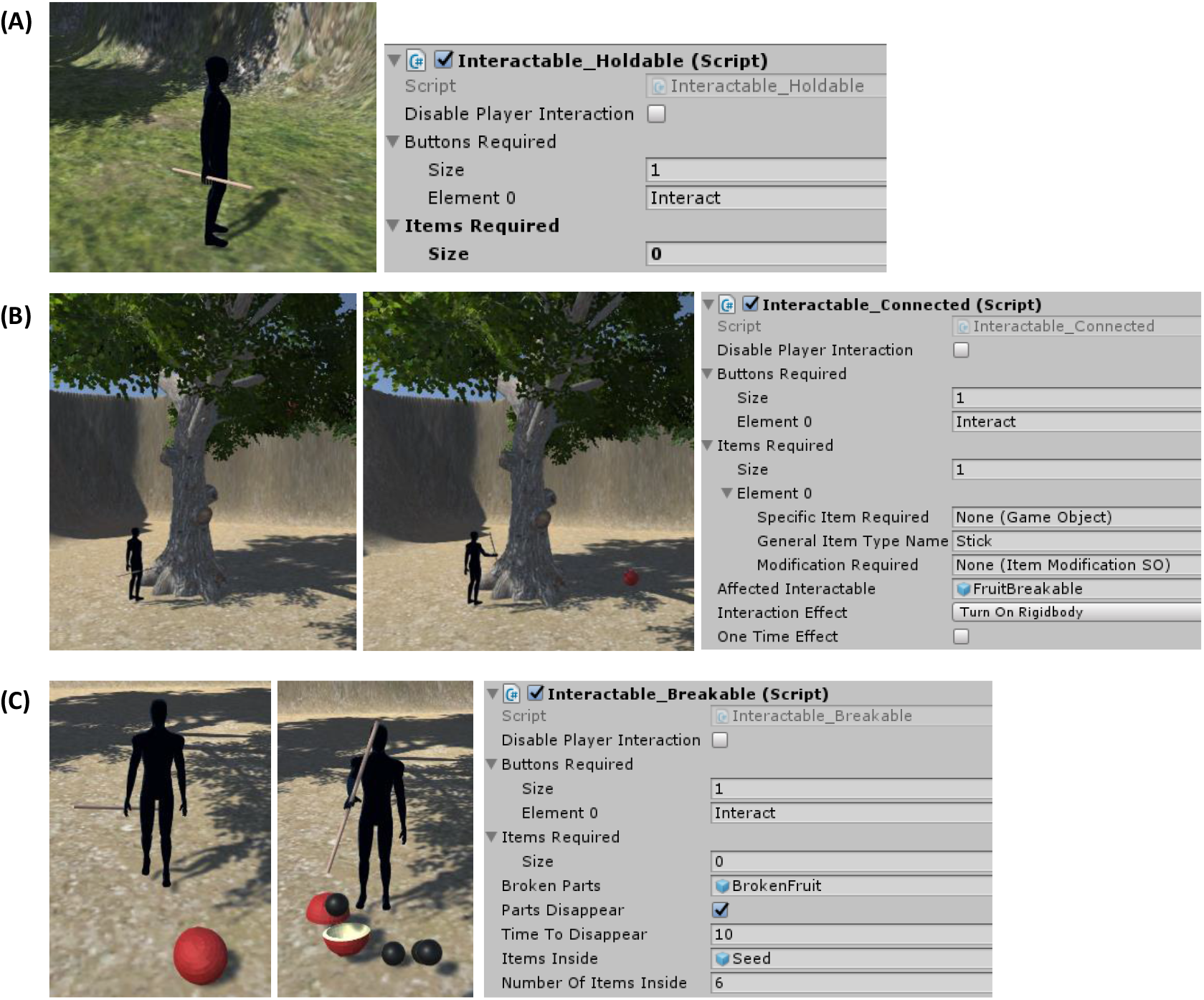
The three main types of `interactable` within the example VERSE environment. Left-hand images show their in-game representations and right-hand images show the particular *Interactable* script attached to the object, along with its properties. **(A)** A `Holdable` ‘stick’ `interactable` which can be picked up by the `player` by pressing the ‘Interact’ key. **(B)** A `Connected` ‘tree’ `interactable` that turns on the Rigidbody component of a connected ‘fruit’ (red ball), thus allowing it to be influenced by gravity and causing it to fall from the top of the tree onto the ground. The tree has one required item for a successful interaction to occur – the `player` must have a ‘stick’ in hand while interacting using the ‘Interact’ key. **(C)** A `Breakable` ‘fruit’ `interactable` which breaks open upon pressing the ‘Interact’ key is when the player is in close vicinity. The fruit becomes a ‘broken fruit’ object (two broken halves) and releases six ‘seeds’ – a `Food interactable` that can be collected by the `player`.

Social information is provided to the participant in the form of two `AIs` that already ‘know’ how to complete the task. Participants are able to observe these `AIs` performing the correct series of behaviours and imitate these behaviours to gain access to the fruit seeds. Both `AIs` contain the following scripts: *AI_Controller, AI_Movement, AI_RandomWalk, AI_ItemInteractionController,* and *AI_FoodInteraction.* For each `AI`, the *AI_RandomWalk* script is programmed so that the `AI` moves around randomly by 2-10 units every 1-5 seconds and is able to detect `Food` during random movement (Figure 4.6A). No ‘attractive areas’ are specified in this example. Movement is permitted via the *AI_Movement* script and is temporarily disabled by the *AI_Controller* script whenever it would interfere with other behaviours such as item interactions.

**Figure 0.6.**
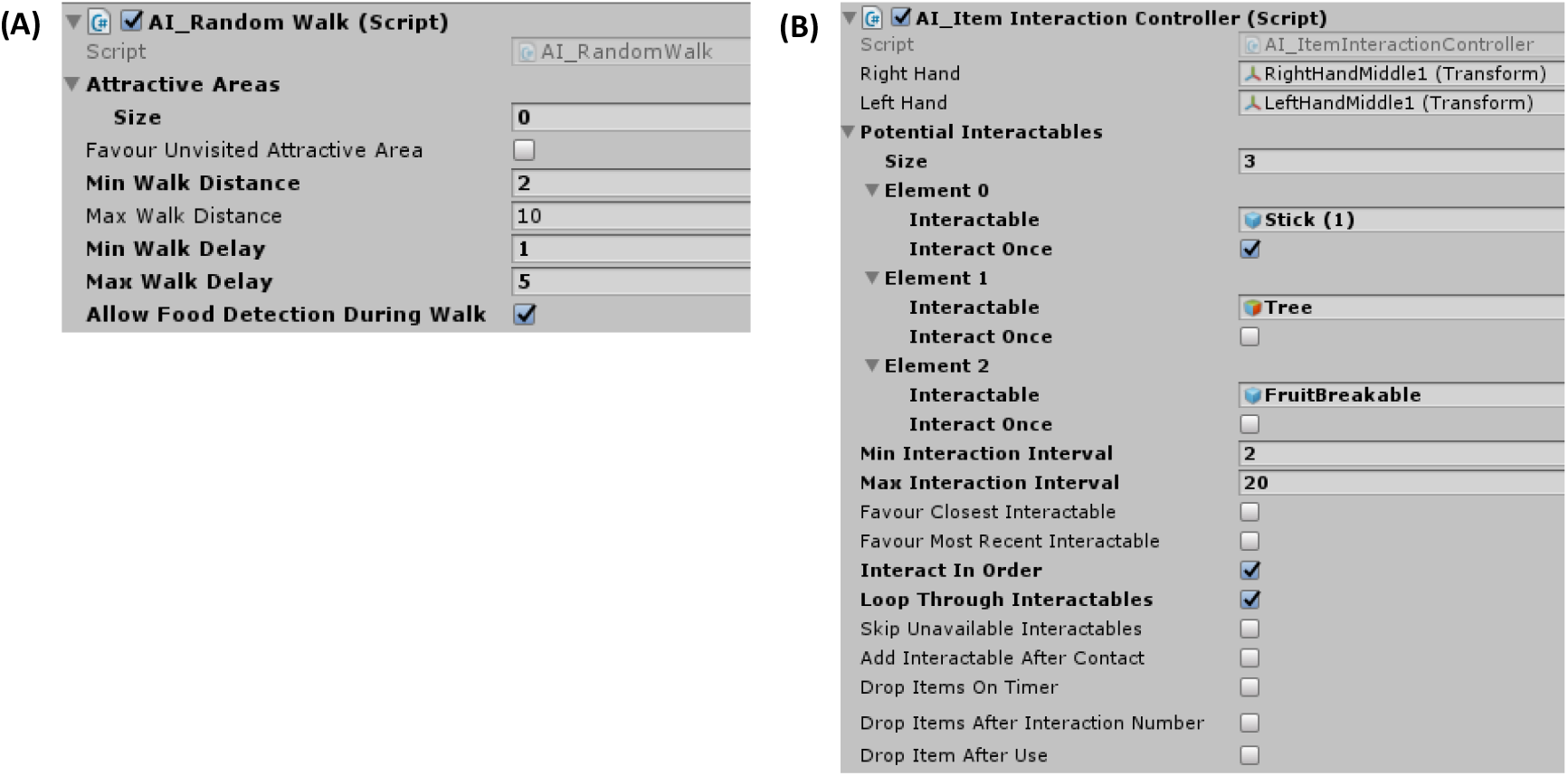
Examples of the *AI_RandomWalk* and *AI_ItemInteractionController* scripts attached to one of the `AIs` in the example environment. **(A)** The *AI_RandomWalk* script, in this example, instructs the `AI` to move a random distance between 2 and 10 units every 1 to 5 seconds and allows `Food` detection during random movement. **(B)** The *AI_ItemInteractionController* script instructs the `AI` to interact with a specified set of `interactables` in a specified way. In this example, the `AI` has the potential `interactables`: ‘Stick (1)’, ‘Tree’ and ‘FruitBreakable’. The `AI` is instructed to interact with the stick object only once, after which it will be removed, and can interact with the other two `interactables` any number of times. The `AI` is instructed to interact with the `interactables` every 2 to 20 seconds in the order they are specified in the list (i.e. the stick first, then the tree, then the fruit) and to loop back to the beginning once it reaches the end of the list. This script can therefore be used to make `AIs` behave in a way that provides social information about cumulative behaviours.

Within the *AI_ItemInteractionController* script (Figure 4.6B), each `AI` has a list of three potential `interactables`. `AIs` are instructed to interact with the `interactables` in this list in order (one interaction every 2-10 seconds) and loop back to the beginning when they reach the end of the list. The ‘potential interactables’ list for each `AI` consists of one of the sticks, one of the trees and the fruit hidden within the branches of the chosen tree. `AIs` are instructed to interact with their chosen stick `interactable` only once, but can interact with the tree and fruit multiple times. Overall, this means that an `AI` will first approach and pick up one of the sticks (after which the stick will be removed from its potential interactables list). It will then approach the tree and interact with it while holding the stick. Since this meets the tree’s interaction requirements, the fruit will fall out of the tree, making it accessible. The `AI` will then approach and break open the fruit, thus gaining access to the seeds inside. The `AI` will then continue to loop through the `interactables` still remaining in its potential interactables list – interacting with the tree, then the fruit, then the tree again, and so on. Since it still has the stick in hand and has not been instructed to drop it, any future interactions with the tree will be successful and cause the fruit to drop. Each interaction occurs after a delay of 2-10 seconds, during which time the `AI`’s other behaviours, such as its random walk, will be in play.

Each `AI` is assigned their own stick, tree and fruit `interactables`. This means that, while both `AIs` will demonstrate to the `player` how to complete the task, each `AI` will provide slightly different sources of social information (i.e. the specific tree used to get the fruit will differ). Finally, the *AI_FoodInteraction* script is set so that each `AI` will detect and collect any `Food` item (in this case, the seeds released from the fruit) within a 2-unit radius and the *AI_RandomWalk* script is programmed to allow `Food` detection during random walks. Thus, when an `AI` breaks open a fruit (or happens to pass a broken fruit while it is moving from one place to another), it will collect any seeds released.

## 6. Discussion and Concluding Remarks

Here I have described a novel experimental tool, VERSE, which takes advantage of gaming technology to allow researchers to create realistic, immersive, open world environments for studying human social learning. VERSE offers a unique way to bring human research in the field of social learning out of the laboratory and into spatially and ecologically relevant scenarios. Where human research has previously lagged behind animal research in the field in terms of ecological validity, this technology offers huge potential for future work on human behavioural ecology and social evolution.

Researchers using VERSE should, however, bear in mind that, while VERSE may offer a more realistic methodology for studying human social behaviour than traditional lab studies, it is still a simplification of real-world social environments. For example, VERSE, in its current form, fails to account for one key component of human social behaviour – verbal communication – which could be an important aspect of our social learning experiences (Rawat, 2016) (although previous research has suggested that children given conflicting information will trust what they see over what they are told; Ma and Ganea, 2010). However, it could be argued, if humans are studied from a behavioural ecology perspective, that *not* including such complex communication gives us a clearer picture of foundational social learning processes (such as stimulus or local enhancement, direct observations and visual cues) that can be analysed in a similar way (and so be directly compared) to studies on animal populations. Indeed, much of the previous work in the field has focussed on how individuals choose to learn based on the decisions of others rather than via direct communications (e.g. Mesoudi, 2008; Morgan *et al.,* 2012; Toelch *et al.,* 2014; Caldwell and Eve, 2014). VERSE could, however, be feasibly extended to include verbal (or written) communication in order to establish its importance in our cultural evolution.

It is also possible, particularly if important aspects of social interaction such as verbal communication are missing, that the artificial intelligence (AI) agents used as social information sources in VERSE may not be viewed by players as truly social entities, but simply as a part of their environment. Evidence from online roleplaying games such as *World of Warcraft* and *Second Life* as to whether non-player characters (NPCs) are viewed as true social interactors is mixed. Often, the identities of NPCs, which generally exist to enrich gameplay and form narratives, are known to the players, which is likely to have a large impact on the extent to which they can be viewed as truly social entities. However, even so, players can form emotional connections with NPCs (Rapp, 2018) and actively collaborate with AI ‘teammates’ (Zhang *et al.,* 2021). Some studies also show that the distinction between NPCs and real players isn’t necessarily clean cut. According to Crenshaw and Nardi (2015), for example, players treated researcher-controlled characters using scripted responses, which could easily be replaced with computer-controlled NPCs, as social cues. Similarly, during Banks and Martey’s (2016) attempts to create ‘transparent’ researcher presence in virtual research environments, a researcher-controlled NPC had to be largely de-humanised to prevent players communicating and interacting with it. In their current form, VERSE AIs are intended to provide pseudo-social cues to establish when and how players copy others – and how this is influenced by the physical form of those social cues. However, further research would be required to establish just how ‘social’ these AI entities are viewed as by players.

It is also important to remember that VERSE is a virtual platform, and this sort of methodology naturally comes with limitations. For example, since VERSE is game-like in its design, participants with different levels of gaming experience may respond differently to this type of methodology – e.g. ‘gamers’ may pick up the computer controls more quickly, thus allowing them to focus their attention on the task at hand more quickly than ‘non-gamers’, who may spend a greater proportion of time learning how to control their player. I would therefore recommend that researchers using VERSE in their experiments do the following to reduce any potential impact of prior gaming ability: (a) allow participants to learn the controls in an initial ‘demo’ environment prior to starting the official experiment, and (b) collect data from each participant concerning prior gaming experience to include in their statistical analysis. Another limitation of VERSE is that participants control a virtual player using computer keys, and hence there is little scope for investigating the learning of motor skills. Despite this, VERSE offers much more flexibility and ecological realism than previous computer-based methods, for example by allowing exploration over large, three-dimensional environments. If future researchers wish to incorporate more realistic motor functions into their experiments, VERSE could feasibly be extended into a fully immersive VR environment, compatible with commercial VR headsets and controllers (e.g. Oculus Quest or HTC Vive), where participants can perform tasks and pick up virtual objects using actual hand movements as opposed to computer keys. In many respects, however, VERSE (or virtual environments in general) offers potential far beyond what is possible in the real world. For example, VERSE could be used to explore human behaviour in scenarios which are impossible to investigate in humans otherwise – such as our responses to dangerous environments, natural disasters and even predators.

Overall, VERSE offers a great deal of potential for researchers wishing to study human behaviour and learning within large, realistic virtual environments. By allowing humans to be studied in spatially explicit environments with the complete behavioural freedom to attempt tasks as they wish, including by observing or ignoring the actions of others, VERSE also offers researchers the opportunity to study humans within a common evolutionary framework alongside animal research. This may even include directly replicating animal experiments using human participants. Due to its modular design, a potentially unlimited number of experiments can be produced in VERSE, including completely novel tasks within completely novel environments, in order to investigate how human social behaviour aids us in realistic survival scenarios. I hope to see future work using VERSE to conduct innovative experiments into human social behaviour.

## Acknowledgements

A big thank you to Will Hoppitt for sparking the idea of creating a VR world for studying human social learning, and to Chris Hassall for your help with structuring the manuscript.

## Data accessibility

VERSE is available at the following figshare repository: https://figshare.com/s/c97c305736c9a3d1c8b9 (Easter, 2022), along with a detailed instruction manual for researchers wishing to use the software. Note that VERSE requires Unity3D to open.

## Footnotes

1 https://unity.com/

2 https://docs.unity3d.com/Manual/GameObjects.html

3 https://docs.unity3d.com/Manual/CreatingAndUsingScripts.html

4 https://docs.unity3d.com/Manual/CreatingScenes.html

5 https://docs.unity3d.com/ScriptReference/Camera.html

6 https://docs.unity3d.com/ScriptReference/Collider.html

7 https://docs.unity3d.com/Manual/class-Rigidbody.html

8 https://docs.unity3d.com/2020.1/Documentation/Manual/UICanvas.html

9 https://docs.unity3d.com/Manual/PublishingBuilds.html

10 https://docs.unity3d.com/Manual/class-ScriptableObject.html

11 https://learn.unity.com/tutorial/inheritance

12 https://assetstore.unity.com

13 https://docs.unity3d.com/2017.4/Documentation/Manual/script-Terrain.html

14 https://docs.unity3d.com/Manual/PhysicsSection.html

15 https://docs.unity3d.com/Manual/class-Rigidbody.html

16 https://docs.unity3d.com/Manual/Navigation.html

17 https://docs.unity3d.com/Manual/Lighting.html

18 https://docs.unity3d.com/Manual/ParticleSystems.html

19 https://assetstore.unity.com/packages/templates/systems/3rd-person-controller-fly-mode-28647

20 https://docs.unity3d.com/ScriptReference/GameObject.SetActive.html

